# A unified model of gene expression control by cohesin and CTCF

**DOI:** 10.1101/2024.11.29.625887

**Authors:** Takeo Narita, Sinan Kilic, Yoshiki Higashijima, Georgios Pappas, Elina Maskey, Chunaram Choudhary

**Affiliations:** Department of Proteomics, The Novo Nordisk Foundation Center for Protein Research, Faculty of Health and Medical Sciences, University of Copenhagen, Blegdamsvej 3B, 2200 Copenhagen, Denmark

## Abstract

Cohesin and CTCF fold vertebrate genomes into loops and topologically associating domains (TADs). The genome folding by cohesin and CTCF is considered crucial for enhancer-promoter communication and gene regulation. However, the extent of cohesin and CTCF-dependent looping in global enhancer function and their potential non-architectural roles in gene activation have remained unclear. Here, we demonstrate that the acute removal of cohesin or CTCF in mouse cells dysregulates hundreds of genes, albeit subtly. Among various possible enhancer types, cohesin almost exclusively facilitates gene activation by one enhancer type—the CBP/p300-dependent enhancers. Interestingly, CTCF plays a dual role, operating both with and without cohesin, and promoting gene activation both in an enhancer-dependent and independent manner. By anchoring cohesin loops, CTCF directs enhancers to specific target genes and prevents their mistargeting. Independently of cohesin and enhancers, CTCF acts as a transcriptional activator or repressor, depending on its precise binding position near promoters. Acting as a canonical transcription activator, CTCF directly activates hundreds of housekeeping genes, including those essential for mammalian cell proliferation. Mechanistically, promoter-bound CTCF controls DNA accessibility and RNA polymerase II recruitment. The transcriptional activator function of CTCF appears unique to vertebrates and is shared by its vertebrate-specific paralog, CTCFL, despite CTCFL’s inability to anchor cohesin loops. These findings reveal the scope of cohesin and CTCF in gene regulation, delineate their shared and unique functions, define a specific class of enhancers using cohesin-mediated looping, and establish a crucial function of CTCF in gene regulation independent of its architectural role. The work reconciles conflicting views and provides a unified model for gene regulation by cohesin and CTCF.

## Introduction

Vertebrate genomes are organized into loops and topologically associating domains (TADs), with cohesin and CTCF playing pivotal roles in this 3D genome organization ^1–5^. Cohesin extrudes loops bidirectionally until it encounters CTCF, where the interaction of cohesin with divergently oriented CTCF molecules halts its processivity ^4^. Cohesin-CTCF-dependent looping facilitates long-range enhancer-promoter (E-P) and promoter-promoter (P-P) interactions. Disrupting CTCF binding sites or inverting their orientation can rewire E-P loops and alter gene expression ^5–9^. The binding of CTCF in promoter-proximal regions directly promotes interactions with distal enhancers ^10–12^ and helps select preferred promoters within a TAD ^13^. In addition, CTCF also functions as a blocker of E-P interactions to prevent enhancer mistargeting ^14–16^. Genome-wide chromatin conformation capture analyses have identified thousands of E-P loops, implying a broad role of 3D genome organization in enhancer-driven gene regulation ^3,11,17–20^. Acute depletion of CTCF or RAD21, an essential cohesin subunit, results in near complete loss of “loop domains.” ^21,22^. However, paradoxically, acute depletion of cohesin or CTCF affects only a limited number of genes ^20–25^, and most E-P interactions are maintained without cohesin or CTCF ^20^.

While the mechanisms of cohesin-CTCF-dependent looping and its broad role in genome folding are well-established, critical questions about their function in global gene regulation remain unanswered. (1) Why does the removal of cohesin or CTCF downregulate only a limited number of genes ^20–25^? (2) Among the multiple types of enhancers in metazoan cells ^26–30^, which enhance types utilize cohesin-dependent looping? (3) Why is there limited overlap in genes affected by cohesin versus CTCF removal ^20,31^? (4) Why is CTCF essential for mammalian cell proliferation but not for *Drosophila* cells, despite its evolutionary conservation ^32–36^? (5) To what extent do cohesin and CTCF regulate expression by non-architectural mechanisms, and what is the nature of such regulated genes?

In this work, we show that the role of cohesin and CTCF in gene regulation is far broader than previously appreciated, though the quantitative impact is subtle. We reveal that cohesin-dependent looping promotes the activation of cell-type-specific genes by a single enhancer type and demonstrates a non-architectural role of promoter-bound CTCF in activating many housekeeping genes. Our findings resolve long-standing debates and provide a unified model of gene regulation by cohesin and CTCF.

## Results

### Strategy for delineating cohesin and CTCF function in enhancer-dependent gene activation

Previous studies have found that only a limited number of genes are affected by the acute depletion of RAD21 and CTCF ^20–25^. However, these studies did not explore why cohesin and CTCF depletion lead to limited transcriptional changes, nor did they investigate whether the affected genes were regulated by enhancers or other mechanisms.

We hypothesized that the roles of cohesin and CTCF in gene regulation could be resolved by first identifying candidate genes regulated by enhancers and then assessing the effects of cohesin and CTCF removal. However, mammalian cells contain diverse types of enhancers that activate genes through distinct coactivators ^27^, and identifying all enhancer types and their specific targets remains challenging.

In our recent work, we found that (1) CBP/p300-catalyzed histone H2B N-terminal acetylation (H2BNTac) preferentially marks active enhancers in mammalian cells ^37^, (2) inhibition of CBP/p300 selectively downregulates genes near candidate enhancers marked by H2BNTac and validated by STARR-seq, while genes lacking these enhancers are largely unaffected ^38,39^, and (3) most CRISPR interference-validated enhancers operate through CBP/p300 ^40^. These findings revealed that CBP/p300 predominantly activates genes via enhancers, and CBP/p300-dependent enhancers constitute a major enhancer type in mammalian cells. Motivated by these findings, in this study we used CBP/p300-dependent enhancers as a prototype to investigate the role of cohesin-CTCF-dependent looping in gene regulation by enhancers. Here, “enhancers” refer specifically to CBP/p300-dependent candidate enhancers, and genes requiring CBP/p300 for activation are considered their putative targets.

By integrating the above findings with prevailing models, we made the following assumptions: (1) cohesin and CTCF regulate transcription through chromatin looping, and (2) CBP/p300 inhibition preferentially impairs transcription of genes activated by enhancers. Therefore, if genes are downregulated after both CBP/p300 inhibition and cohesin/CTCF depletion, this indicates that they require both chromatin looping and enhancers for activation. Conversely, if genes are downregulated by cohesin/CTCF depletion but not by CBP/p300 inhibition, they depend on chromatin looping but are activated without enhancers or require enhancers that function without CBP/p300. Finally, if genes are downregulated by CBP/p300 inhibition but not by cohesin/CTCF depletion, they are possibly activated by enhancers without requiring cohesin-CTCF-dependent looping.

### Cohesin and CTCF regulate hundreds of genes but with quantitatively modest effects

To define cohesin- and CTCF-regulated genes, we used mouse embryonic stem cells (mESCs) expressing degron-fused *Rad21-mAid-Gfp* (RAD21^AID^) and *Ctcf-mAid-Gfp* (CTCF^AID^) ^22,23^. RAD21^AID^ and CTCF^AID^ are rapidly and effectively depleted in these cells ^22,23^. RAD21^AID^ depletion impairs cell proliferation more rapidly than CTCF^AID^ depletion ^22,23^, so we depleted RAD21^AID^ for 4 hours and CTCF^AID^ for 6 hours, then quantified transcriptional changes in two biological replicates using 5-ethynyl uridine RNA labeling and next-generation sequencing (EU-seq). CBP/p300-regulated genes were identified by treating cells with the CBP/p300 inhibitor A-485 for 0.5–2 hours ^39^. The specificity of A-485 was confirmed by comparisons with CBP/p300 genetic deletion and PROTAC-induced depletion ^38,41^.

CBP/p300 inhibition significantly downregulated over 1,000 genes (P_adj_ < 0.05), while RAD21^AID^ and CTCF^AID^ depletion depletion affected only 1 and 37 genes, respectively (**Supplemental Figure 1A**). The limited transcription changes by CTCF and RAD21 are consistent with a recent study showing that only 55 (out of ∼30,000 genes analyzed) genes are significantly regulated after acute (3h) RAD21^AID^ and CTCF^AID^ depletion ^20^.

We hypothesized that gene expression changes following RAD21^AID^ and CTCF^AID^ depletion might be too subtle to meet statistical significance. To capture weakly regulated genes, we used a fold-change-based criterion, categorizing genes based on their degree of regulation: highly down- or up-regulated (HD or HU; ≥2-fold change), intermediate down- or up-regulated (ID or IU; 1.5–2-fold change), slightly down- or up-regulated (SD or SU; 1.3–1.5-fold change), and not changed (NC; <1.2-fold change) (**Supplemental Figure 1B**). A-485 regulated genes were classified as very highly downregulated (HD2; ≥4-fold decrease), highly downregulated (HD1; ≥2-fold decrease), intermediate downregulated (ID; ≥1.5-2-fold decrease), and not downregulated (ND; <1.2 decrease). Genes with low expression (TPM <15) were excluded to minimize background noise, resulting in ∼9,500 genes quantified per perturbation. This analysis showed that RAD21^AID^ and CTCF^AID^ depletion affect several hundred genes, though the changes are modest (**Supplemental Figure 1B**).

**Supplemental Figure 1.**
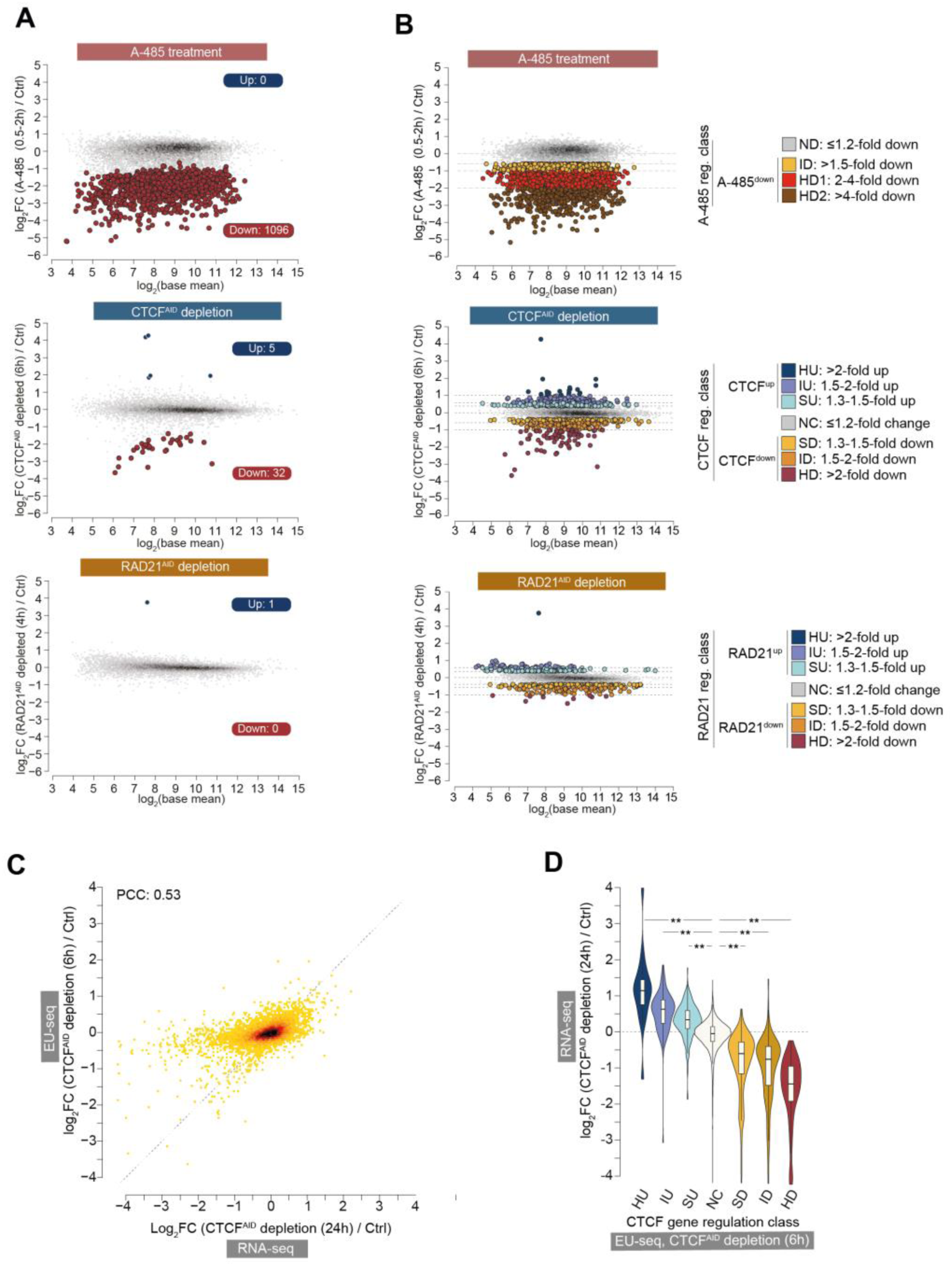
The scope of CTCF, RAD21, and CBP/p300-dependent enhancers in gene regulation and classification of regulated genes, related to Figure 1. **(A)** The number of significantly (Padj <0.05) up- and down-regulated genes after acute CTCF^AID^ depletion (6h), RAD21^AID^ (4h) depletion, and A-485 treatment (0.5-2h). Transcription changes were quantified in mESCs expressing CTCF^AID^ and RAD21^AID^ using 5-ethylidine uridine nascent RNA labeling and next-generation sequencing (EU-seq) (n =2). Transcription changes are compared in cells without or with depletion of CTCF^AID^ and RAD21^AID^ by treatment with IAA (500uM). A-485-induced nascent transcription data are from reference ^39^. The analysis includes all genes expressed at TPM >5. **(B)** Fold-change-based classification of genes regulated after acute CTCF^AID^ (6h) depletion, RAD21^AID^ (4h) depletion, and A-485 treatment (0.5-2h). Regulated genes are classified using the indicated fold-change in gene expression after the specified treatments. The analysis only includes genes that are expressed at TPM >15. **(C)** Correlation between gene regulation after acute (6h) and long-term (24h) depletion of CTCF^AID^. Transcription changes were quantified by EU-seq after acute CTCF^AID^ depletion (this study), and by RNA-seq after long-term CTCF^AID^ depletion. RNA-seq data are from reference ^42^. Pearson correlation coefficient (PCC) is indicated. **(D)** Change in expression of acutely regulated CTCF target genes after long-term CTCF^AID^ depletion. After acute (6h) CTCF^AID^ depletion, up- and down-regulated genes are classified into the indicated class, as specified in Supplemental Figure1B. Within each class, change in gene expression after long-term CTCF depletion (24h, RNA-seq) is shown. RNA-seq data are from reference ^42^. The box plots display the median, upper and lower quartiles; the whiskers show the 1.5× interquartile range (IQR). Two-sided Mann–Whitney U-test, followed by correction for multiple comparisons with the Benjamini–Hochberg method; **Padj<0.001.

### Modest gene expression changes reflect genuine biological regulation

Small fold changes may indicate genuine regulation or arise from quantification noise. To assess reliability, we compared our CTCF-regulated genes with published RNA-seq data from the same CTCF^AID^ cells. Although the overall correlation was modest (PCC: 0.53), genes identified by EU-seq fold change were significantly (Padj; HU versus NC 7.1e-11, IU versus NC 2.3e-40, SU versus NC 2.8e-52, SD versus NC 9.2e-62, ID versus NC 1.5e-46, HD versus NC 8.4e-44, Two-sided Mann–Whitney U-test) regulated in RNA-seq (**Supplemental Figure 1C-D**), even though the experiments were conducted at different times (6h vs. 24h) and by different labs. This suggests our fold-change-based approach likely captures primary CTCF targets.

RAD21-regulated genes showed smaller fold changes than CTCF-regulated genes (**Supplemental Figure 1B**). To test the reliability of RAD21-regulated genes, we conducted 12 additional EU-seq replicates in RAD21^AID^ cells. In these new data, 20 genes were significantly downregulated (Padj < 0.05) and 1 upregulated (**Supplemental Figure 2A**). While more replicates increased the number of significantly regulated genes, most genes with <2-fold changes remained below statistical thresholds.

We next examined fold-change reproducibility, analyzing genes from initial experiments and measuring the fraction downregulated by ≥1.1-, 1.2-, or 1.3-fold in the new replicates. Overall, 69.5% of genes downregulated in initial experiments were also downregulated by ≥1.3-fold in at least half of the new replicates, whereas only 1.3% of genes not downregulated initially showed ≥1.3-fold downregulation in the new replicates (**Supplemental Figure 2B**). Among initially downregulated genes, over 90% of HD+ID genes and 62% of SD genes remained downregulated by ≥1.3-fold in at least half (6/12) of the new replicates. True reproducibility is likely even higher, as some genes initially downregulated fell below the 1.3-fold threshold in new replicates. For further analyses, we included only genes downregulated by ≥1.3-fold in the initial RAD21^AID^ depletion and by ≥1.3-fold in at least half of the new replicates (**Supplemental Figure 2C**). Together, this empirical analysis and additional analyses (see below, **Supplemental Note 1**) support the reliability of our fold-change-based approach.

**Supplemental Figure 2.**
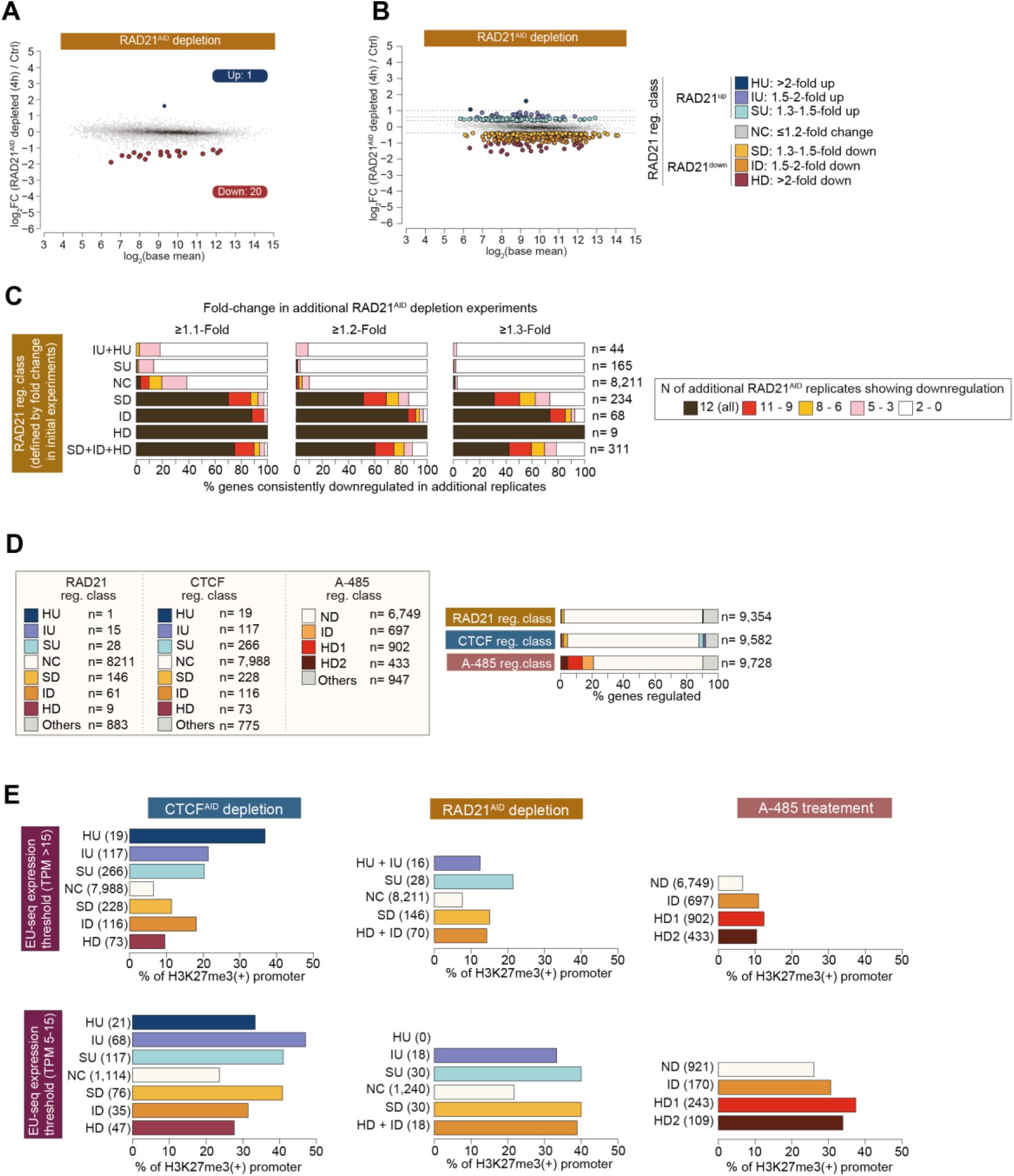
Early CTCF targets are consistently regulated after long-term CTCF depletion, and most genes regulated by cohesin and CTCF are not repressed by polycomb, related to Figure 1. **(A)** The number of significantly (Padj <0.05) up- and down-regulated genes in mESC after acute (4h) RAD21^AID^ depletion. Transcription changes were quantified using EU-seq (n=12). The analysis includes all genes expressed at TPM >5. The gene expression threshold is determined from EU-seq expression values under the control condition. **(B)** Genes up- and down-regulated by RAD21^AID^ depletion (n= 12 replicates). Up- and down-regulated genes are grouped into indicated gene regulation classes. The analysis only includes genes that are expressed at TPM >15. The gene expression threshold is determined from EU-seq expression values under the control condition. **(C)** Reproducibility of gene regulation by RAD21^AID^ depletion in mESC. RAD21-regulated genes were classified based on fold changes in initial experiments (n =2). Subsequently, gene expression changes were quantified in 12 additional replicates. Within the RAD21-regulated gene class defined from initial experiments, shown is the fraction of genes showing ≥1.1, ≥1.2, and ≥1.3 fold-change in the indicated number of additional replicates. **(D)** Shown is the number of genes in the indicated gene regulation class. Gene regulation class is defined based on fold-change after RAD21^AID^ depletion, CTCF^AID^ depletion, and A-485 treatment. For CTCF^AID^ depletion and A-485 treatment, genes are classified based on data from two biological replicates. For RAD21^AID^ depletion, genes are classified based on 14 replicates (2 initial replicates and 12 additional replicates). Fraction of genes regulated after RAD21^AID^ depletion, CTCF^AID^ depletion, and A-485 treatment (right panel). **(E)** Genes downregulated by CTCF^AID^ depletion, RAD21^AID^ depletion, and A-485 treatment show minimal difference in polycomb-catalyzed H3K27me3 enrichment. Genes regulated by the specified treatments are grouped into high (TPM >15) and low (TPM 5-15) expressed, and classified into the indicated regulated gene class, as defined in Supplemental Figure1B. Within the specified conditions, and regulated gene class, genes exhibiting H3K27me3 in promoter regions (+/-2kb from TSS) is shown.

### Cohesin-dependent looping specifically facilitates gene activation by CBP/p300

For the remainder of the study, we used a fold-change-based approach to define regulated genes (**Supplemental Figure 2D**). Gene regulation categories for each treatment condition are noted in superscript text. For instance, genes downregulated following RAD21^AID^ and CTCF^AID^ depletion are labeled as RAD21^down^ and CTCF^down^, respectively, while upregulated genes are labeled RAD21^up^ and CTCF^up^. Unchanged genes are denoted as RAD21^NC^ and CTCF^NC^. A-485 downregulated and not downregulated genes are referred to as A-485^down^ and A-485^ND^, respectively.

RAD21- and CTCF-regulated genes display distinct regulation by CBP/p300 inhibition (**Figure 1A**). For example, RAD21^down^ genes are strongly biased for A-485^down^ genes, while RAD21^up^ genes show no such bias. Notably, within RAD21^down^ genes, those most strongly affected by RAD21^AID^ depletion also show significantly stronger downregulation with A-485 treatment (**Figure 1B**). CTCF-regulated genes show an opposite trend: CTCF^up^ genes are more strongly biased for A-485^down^ genes than CTCF^down^ genes, and among CTCF^down^ genes, only weakly downregulated show stronger downregulation by A-485 (**Figure 1A-B**).

**Figure 1.**
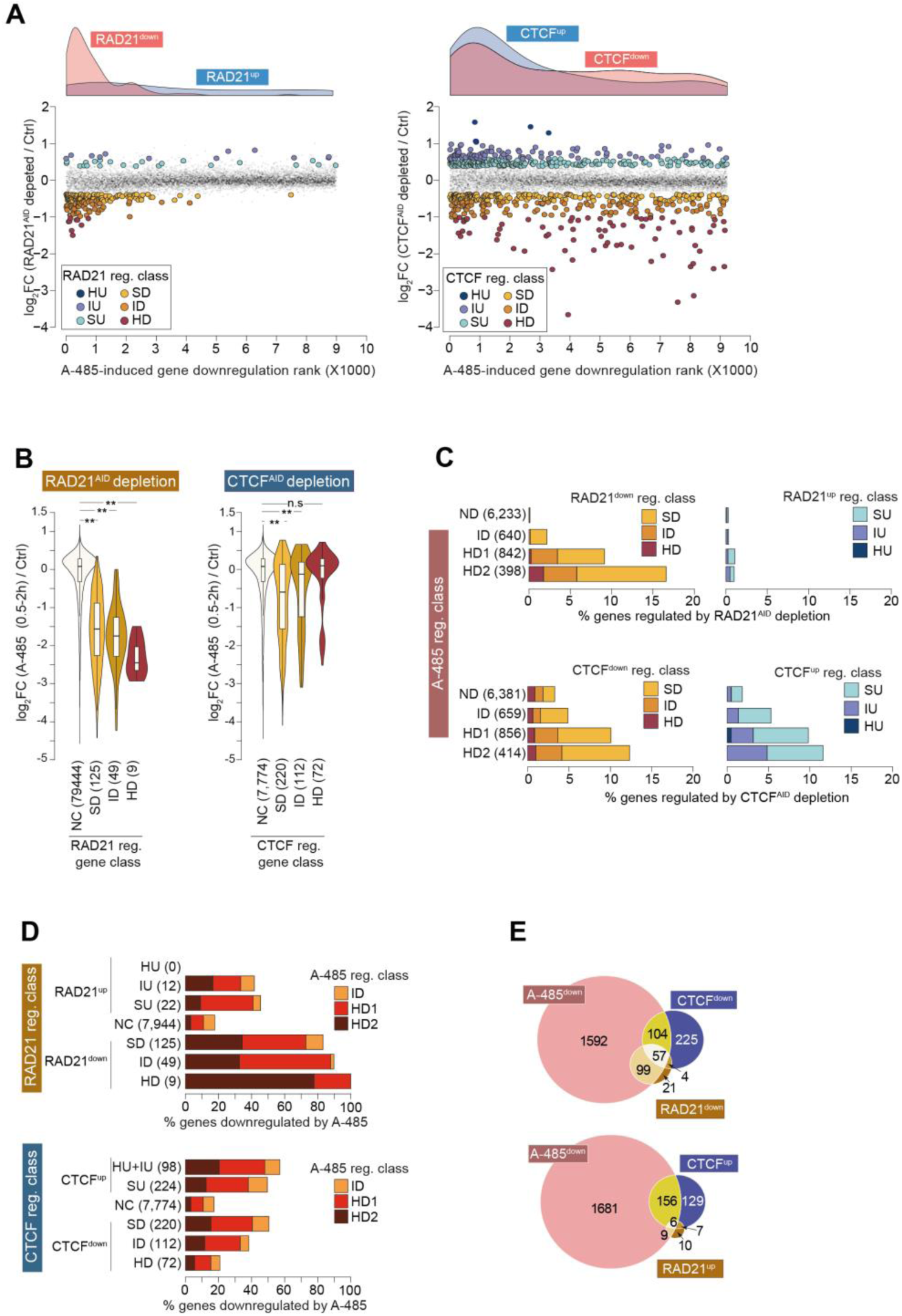
Cohesin and CTCF regulate hundreds of genes, but subtly. **(A)** Fold-change in gene expression after acute depletion of RAD21^AID^ (4h, left panel) and CTCF^AID^ (6h, right panel). Genes are rank-ordered by the extent of their downregulation after A-485 treatment, and the density plots (top) show the distribution of genes up- and down-regulated after RAD21^AID^ and CTCF^AID^ depletion. Genes up- and down-regulated after RAD21^AID^ and CTCF^AID^ depletion are classified as specified in Supplemental Figure 1B. **(B)** A-485-induced regulation among different groups of genes regulated by RAD21^AID^ and CTCF^AID^ depletion. The box plots display the median, upper and lower quartiles; the whiskers show the 1.5× interquartile range (IQR). Two-sided Mann–Whitney U-test, followed by correction for multiple comparisons with the Benjamini–Hochberg method; **Padj<0.001, *Padj<0.05. **(C)** Fraction of RAD21 and CTCF up- and down-regulated genes within different groups of A-485 regulated genes. A-485 regulated genes are grouped into the indicated categories, as defined in Supplemental Figure 1B. Within each category, the fraction of genes upregulated or downregulated after acute depletion of RAD21^AID^ (top panels) and CTCF^AID^ (bottom panels) is depicted. **(D)** Fraction of A-485-downregulated genes within different RAD21 and CTCF-regulated gene groups. Genes regulated after CTCF^AID^ and RAD21^AID^ depletion are grouped into the indicated classes. Within each class, the fraction of genes regulated by A-485 is shown. **(E)** Overlap of CTCF^down^ and RAD21^down^ genes (top panel) and CTCF^up^ and RAD21^up^ genes (bottom panel) with A-485-downregulated genes. See also Supplemental Figure 1.

In the RAD21^down^ gene sub-categories, the fraction of RAD21^down^ genes increases with the degree of downregulation by A-485, while RAD21^up^ genes show minimal difference (**Figure 1C**). Notably, RAD21^down^ genes constitute 16.6% of the HD2 category, a 127-fold increase over the ND category (0.13%; odds ratio 199, p < 2.2e-16, Fisher’s exact test).

CTCF can either facilitate or block E-P interactions ^14,15,30,43^. Consistently, both CTCF^down^ and CTCF^up^ genes are biased for regulation by CBP/p300, and the fraction of CTCF^down^ and CTCF^up^ genes increase similarly with an increasing magnitude of A-485-induced gene downregulation (**Figure 1C**). This implies that CTCF’s function in facilitating CBP/p300-dependent enhancer targeting and blocking enhancer mistargeting might be similarly broad.

Among RAD21^down^ genes, the fraction of A-485-downregulated genes increases with increasing magnitude of RAD21^AID^ depletion-induced gene downregulation (SD, 83%; ID, 90%; HD, 100%) (**Figure 1D**). CTCF^down^ genes exhibit the opposite trend; as the magnitude of CTCF^AID^ depletion-induced gene downregulation increases, the fraction downregulated by A-485 declines (SD: 50%, ID: 38%, HD: 21%). Conversely, as the magnitude of CTCF^AID^ depletion-induced gene upregulation increases, the fraction of CTCF^up^ genes overlapping with A-485^down^ genes increases.

Genes regulated by RAD21 and CTCF exhibit only partial overlap, and the overlapping and non-overlapping genes show differences in their overlap with CBP/p300-regulated genes (**Figure 1E**). A majority (86%) of RAD21^down^ genes overlap with A-485^down^ genes (odds ratio 80, p < 2.2e-16), whereas only 44.1% of RAD21^up^ genes do (odds ratio 3.8, p = 3.0e-4). Conversely, 41.8% of CTCF^down^ genes overlap with A-485^down^ genes (odds ratio 3.4, p < 2.2e-16), and 51.9% of CTCF^up^ genes overlap (odds ratio 6.0, p < 2.2e-16). RAD21^down^ genes constitute only 2.2% of genes quantified in mESC, but 32.2% of RAD21^down^ genes overlap with CTCF^down^ genes (odds ratio 17, p < 2.2e-16). Among the commonly downregulated genes by RAD21^AID^ and CTCF ^AID^ depletion, 93.4% overlap with A-485^down^ genes (odds ratio 71, p < 2.2e-16).

In addition to regulating enhancer-promoter interactions, cohesin also counters interactions between polycomb domains, and depletion of RAD21 leads to repression of polycomb targets ^23^. In our analyses, the early gene downregulation following RAD21^AID^ depletion appears to primarily result from diminished activation by CBP/p300-dependent enhancers rather than polycomb-mediated repression (**Supplemental Figure 2E**, **Supplemental Note 2**).

Together, these results demonstrate that: (1) cohesin primarily promotes gene activation by CBP/p300-dependent cis-regulatory elements. (2) CTCF also promotes gene activation by CBP/p300-dependent enhancers, but most CTCF-dependent genes are activated without CBP/p300. (3) Contrary to expectations, genes strongly downregulated by CTCF^AID^ depletion are less likely to be activated by CBP/p300 than those more weakly downregulated.

### Genes downregulated by RAD21^AID^ depletion reside in enhancer-rich regions

Next, we examined the prevalence of candidate enhancers near regulated genes. Enhancers were identified by the presence of H2BK20ac, a constituent of the H2BNTac signature ^37^, and their activity was gauged by H2BK20ac ChIP signal strength. Unlike RAD21^down^ genes, most CTCF^down^ genes do not overlap with A-485^down^genes (**Figure 1E**). Thus, we separated CTCF^down^ genes into two groups: those downregulated by both CTCF^AID^ depletion and A-485 treatment (CTCF^down^A-485^down^) and those downregulated by CTCF^AID^ depletion alone (CTCF^down^A-485^ND^).

RAD21^down^ genes, unlike randomly selected genes, show a notably stronger enrichment of H2BK20ac-positive enhancers in their vicinity. Similarly, CTCF^down^A-485^down^ genes are enriched with proximal enhancers, while CTCF^down^A-485^ND^ genes are not (**Figure 2A**). CTCF^down^A-485^down^ genes show a tendency of downregulated by RAD21^AID^ depletion, whereas CTCF^down^A-485^ND^ genes do not. This absence of nearby enhancers may explain why CTCF^down^A-485^ND^ genes are not affected by CBP/p300.

**Figure 2.**
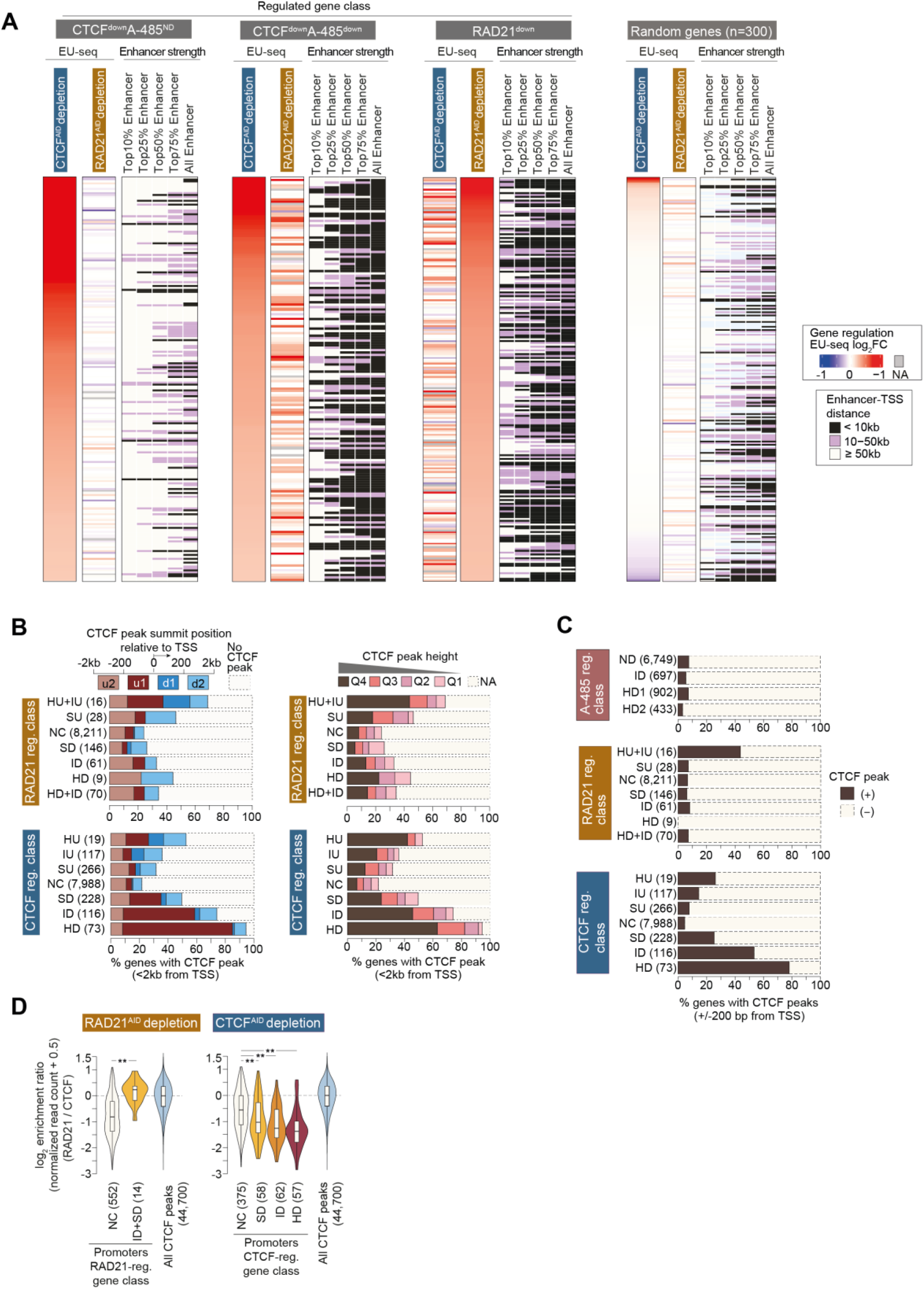
Genes activated by CTCF exhibit strong CTCF binding in their promoters. **(A)** Candidate enhancer enrichment in proximity to genes downregulated after RAD21^AID^ and CTCF^AID^ depletion and randomly selected genes in mESC. CTCF downregulated genes are grouped into two categories: CTCF^down^A-485^down^ and CTCF^down^A-485^ND^. Enhancers are defined by H3K27ac and H2BK20ac overlapping peaks, excluding region +/-500bp from the TSS, and enhancer strength is determined by H2BK20ac ChIP signal enrichment. **(B)** CTCF binding in promoter-proximal (+/- 2kb) regions. Genes regulated after CTCF^AID^ and RAD21^AID^ depletion are grouped into the specified classes. Within each class, the fraction of genes having CTCF peaks within the indicated distance from the TSS is displayed (left panel). The right panel shows CTCF peak strength within the indicated classes of regulated genes. **(C)** Enrichment of CTCF near promoters. Genes regulated after A-485 treatment, CTCF^AID^ depletion, and RAD21^AID^ depletion were grouped into the specified classes. The fraction of genes with CTCF peaks within +/-200bp from their TSS is depicted within each class. **(D)** Shown is RAD21/CTCF enrichment ratio in all CTCF peaks, and CTCF peaks found in the promoters of the indicated classes of RAD21 and CTCF-regulated genes. Only the genes bound by both CTCF and RAD21, within +/- 200bp from their TSS, are included in this analysis. The box plots display the median, upper and lower quartiles; the whiskers show the 1.5× interquartile range (IQR). Two-sided Mann–Whitney U-test, followed by correction for multiple comparisons with the Benjamini–Hochberg method; **Padj<0.001, *Padj<0.05. See also Supplemental Figures 1 and 2.

**Supplemental Figure 3.**
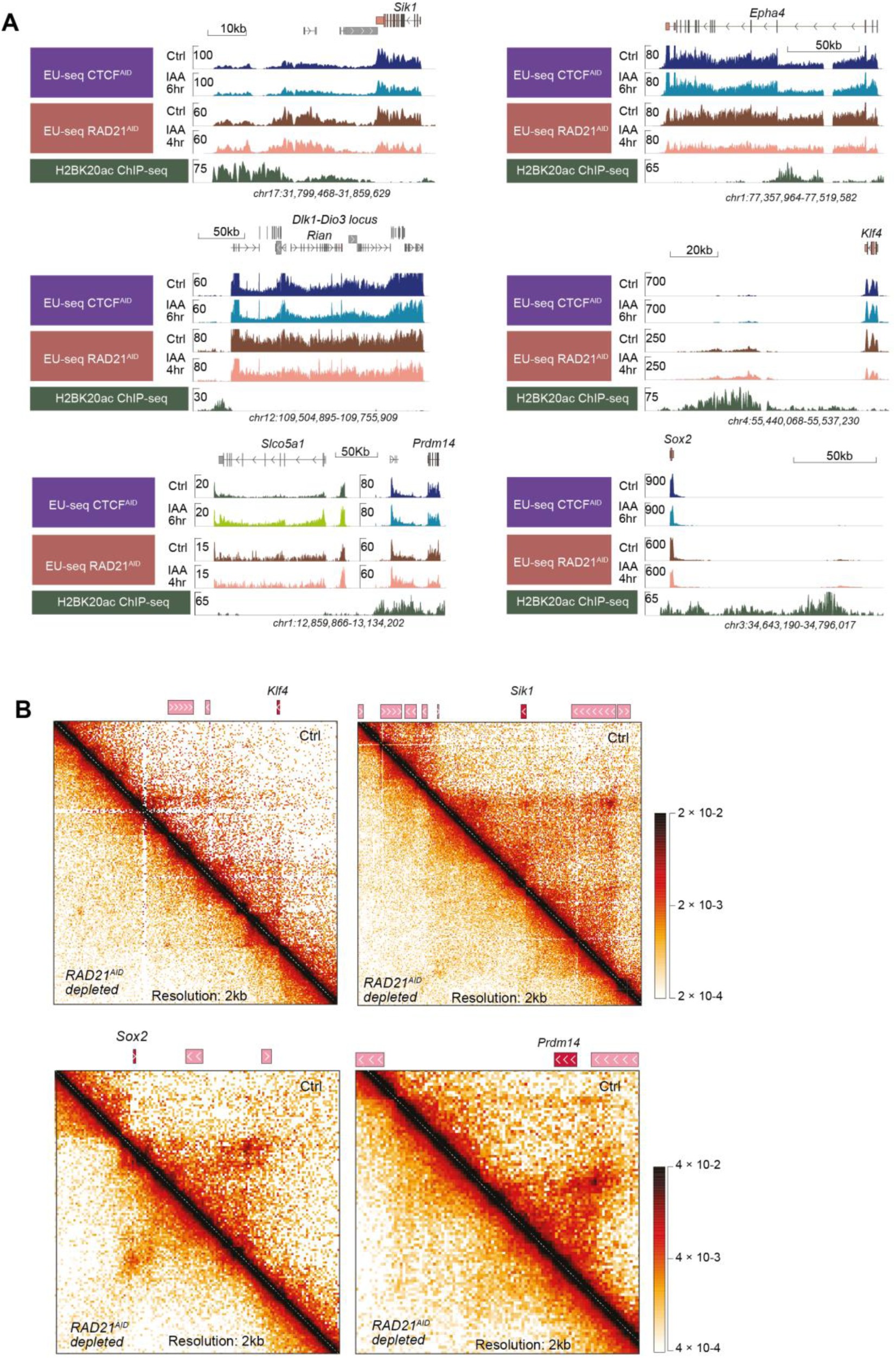
RAD21^AID^ and CTCF^AID^ depletion decrease the expression of known enhancer targets, and RAD21^AID^ depletion causes loss of chromatin loops in the regulated gene loci, related to Figure 1 and 2. **(A)** Genome browser tracks showing downregulation of known enhancer target genes by RAD21^AID^ and CTCF^AID^ depletion in mESC. H2BK20ac ChIP tracks show the position of known enhancers for the indicated genes. Of note, *Sik1*, *Klf4*, and *Sox2* show variable levels of nascent transcription in non-genic regions marked by H2BK20a(C) but this transcription in non-genic regions is not reduced by RAD21^AID^ and CTCF^AID^ depletion, whereas transcription of the known target genes is reduced. *Klf4* expression is only reduced after RAD21^AID^ depletion, and expression of Slco5a1 is specifically increased after CTCF^AID^ depletion, consistent with re-targeting of enhancer to this gene after removal of CTCF binding in this locus ^46^. **(B)** Micro-C detected chromatin interactions in the indicated enhancer target loci in wild-type mESC, and weakening of chromatin loops after RAD21^AID^ depletion. Micro-C data are from reference ^20^.

**Supplemental Figure 4.**
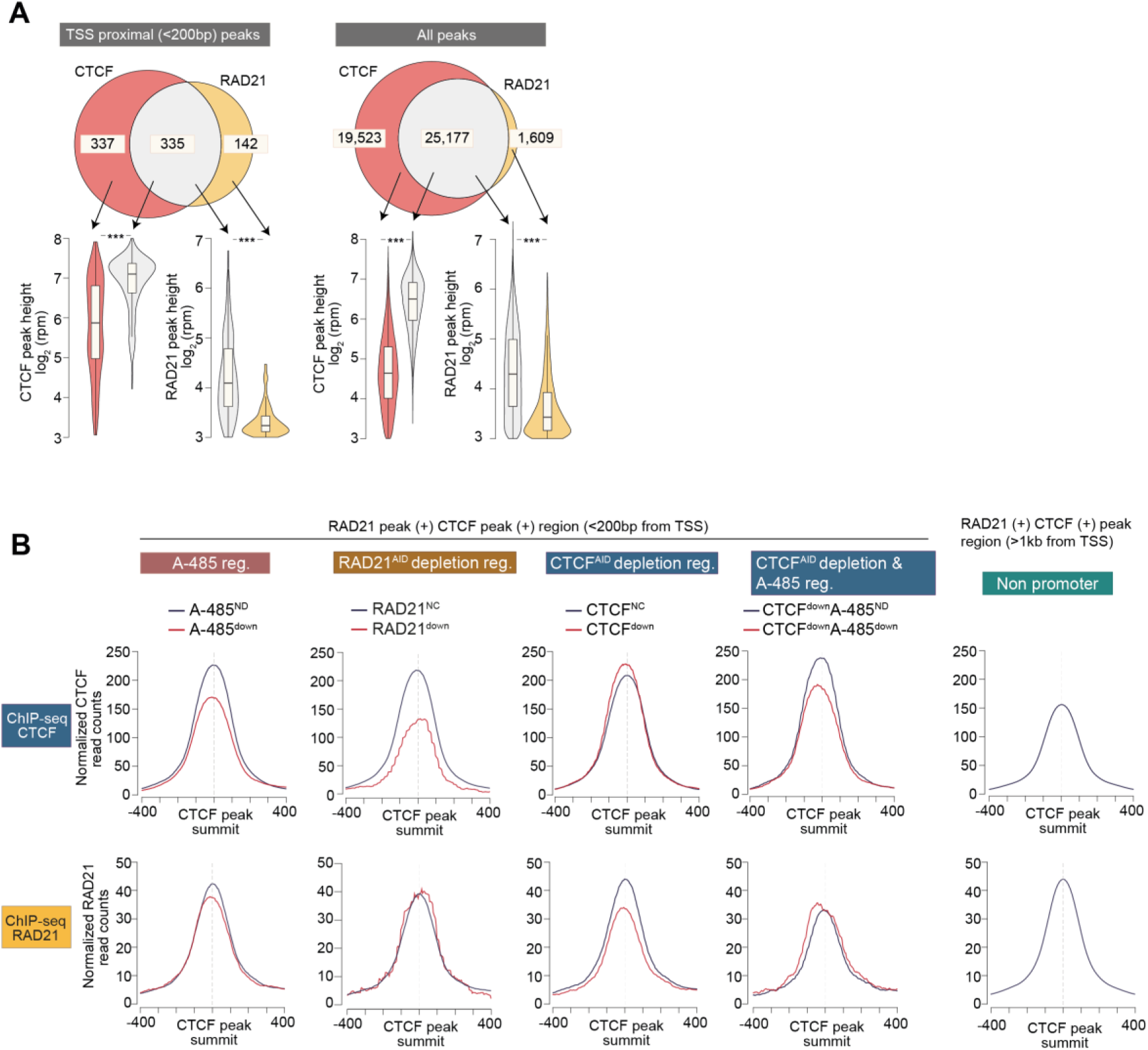
CTCF^down^ genes show relatively high enrichment of CTCF and poor enrichment of RAD21 in their promoters, related to Figure 3. **(A)** The top panels show overlap between CTCF and RAD21 ChIP-seq peaks detected in TSS proximal (+/-200bp from TSS) regions and all regions. One RAD21 peak can overlap with more than one CTCF peak or vice versa, and the higher number of overlapping peaks is shown. The bottom panels show relative enrichment of CTCF and RAD21 in TSS-proximal and distal peaks. The box plots display the median, upper and lower quartiles; the whiskers show the 1.5× interquartile range (IQR). Two-sided Mann–Whitney U-test, followed by correction for multiple comparisons with the Benjamini–Hochberg method; **Padj<0.001. Of note, CTCF peak height is similar, or even higher in TSS proximal peaks than in distal peaks, but RAD21 shows lower enrichment in TSS proximal regions as compared to distal regions. **(B)** Aggregate plots showing CTCF and RAD21 enrichment in the indicated groups of promoters and non-promoter regions. This analysis only includes promoters that are bound (within +/- 200bp of TSS) by CTCF and RAD21 in the analyzed ChIP-seq data. Promoters are grouped based on the regulation of corresponding genes after the specified treatments. Peaks are centered using CTCF peak summit.

To confirm that RAD21^down^ and CTCF^down^ genes include those regulated by enhancers, we analyzed several genes known to depend on enhancers and/or cohesin-mediated looping, including *Sox2*, *Klf4*, *Epha4*, *Sik1*, *Dlk1*-*Dio3*, and *Prdm14*. All were downregulated by both RAD21^AID^ and CTCF^AID^ depletion, except Klf4, which was affected only by RAD21^AID^ depletion (**Supplemental Figure 3A**). At the *Prdm14* locus, loss of a CTCF-anchored loop leads to enhancer retargeting from *Prdm14* to *Slco5a1*, with *Prdm14* expression reduced in both CTCF^AID^ and RAD21^AID^ depleted cells, but *Slco5a1* expression increasing only upon CTCF^AID^ depletion. This suggests that enhancer redirection in CTCF^AID^-depleted cells results not from a simple loss of looping; instead, re-targeting occurs due to the failure to halt cohesin-mediated looping. Thus, cohesin is still required for the re-targeting to occur. Supporting the looping model, Micro-C data confirm cohesin-dependent loops in the loci of *Sox2, Sik1, Klf4,* and *Prdm14* (**Supplemental Figure 3B**). Regulation of most of these genuine enhancer targets will be overlooked in standard analyses due to statistical thresholds. Together, these data support the role of cohesin-CTCF-dependent looping in enhancer-dependent gene activation.

### Genes downregulated by CTCF^AID^ depletion are bound by CTCF binding in their promoters

CTCF binding near gene promoters can interactions with distal enhancers ^10–12^. CTCF binds more frequently within ±2 kb of CTCF-regulated genes compared to non-regulated genes, with its enrichment correlating to the level of gene regulation (**Figure 2B**). Notably, in CTCF^down^ genes, CTCF binds very close to promoters. Promoter CTCF binding is suggested to guide enhancer target selection within a domain ^13^. We examined CTCF binding in promoters (within ±200bp of TSS) to test if this is a general principle. Both CTCF^down^ and CTCF^up^ genes exhibit frequent promoter-proximal CTCF binding, although CTCF^down^ genes show a higher prevalence. This binding increases with the extent of gene regulation upon CTCF^AID^ depletion (**Figure 2C**). In contrast, CTCF is present in the promoters of <8% of A-485^down^ and RAD21^down^ genes, with no correlation to the magnitude of gene downregulation following A-485 treatment or RAD21^AID^ depletion. Together, these results confirm that CTCF is enriched in the promoters of genes downregulated by CTCF depletion ^20,22,25,31,44,45^. However, most RAD21^down^ genes lack CTCF binding in their promoter, suggesting that promoter CTCF binding is not a general mechanism for directly guiding enhancers to their target genes.

To explore the function of CTCF at the promoters of CTCF^down^ genes, we assessed whether it might facilitate cohesin-mediated interactions with enhancers that do not rely on CBP/p300. If so, CTCF-bound promoters should show a co-enrichment of cohesin. We compared the RAD21/CTCF enrichment ratio at promoters versus all CTCF peaks. In RAD21^down^ gene promoters, the RAD21/CTCF ratio resembles that of distal CTCF peaks, though the number of such genes is limited (**Figure 2D**, **Supplemental Figure 4A-B**). In CTCF^down^ gene promoters, however, the RAD21/CTCF ratio is significantly lower than at distal CTCF peaks and declines with increased downregulation following CTCF^AID^ depletion. This indicates that CTCF binding in these CTCF^down^ gene promoters is not prominently involved in halting cohesin loops.

### Promoter-bound CTCF activates genes independently of anchoring cohesin loops

Mutating CTCF at Y226A and F228A selectively disrupts its ability to anchor cohesin loops ^47,48^. To delinate cohesin-dependent and independent functions of CTCF, we engineered mESCs to express a degron-tagged mutant CTCF (*Gfp-Fkbp12^V36F^-Ctcf^Y22A^*^6^*^/F^*^228^*^A^*; hereafter CTCF^YF-AA^) (**Supplemental Figure 5A**). Treatment with dTAG-13 rapidly degraded CTCF^YF-AA^, confirming efficient depletion (**Supplemental Figure 5B-C**).

Like human HAP1 cells expressing CTCF^YF-AA^, mESCs with CTCF^YF-AA^ continued to proliferate, and CTCF^YF-AA^ depletion impaired growth (**Supplemental Figure 5D**). Nonetheless, cells expressing CTCF^YF-AA^ showed marked transcriptional alterations compared to control CTCF^AID^ expressing cells, with similar numbers of genes showing up- and down-regulation (**Supplemental Figure 5E**). The dysregulated genes show a bias toward A-485^down^ genes (**Supplemental Figure 5F-G**), highlighting a probable role for CTCF-dependent cohesin loop anchoring in enhancer-dependent gene regulation. However, these changes reflect both primary and secondary effects, making it challenging to isolate direct impacts.

To identify CTCF functions retained by CTCF^YF-AA^, we measured transcriptional changes after acute depletion of CTCF^YF-AA^ (2–4 hours). Within 2 hours, hundreds of genes showed reduced expression, including many downregulated by wild-type CTCF^AID^ depletion (**Supplemental Figure 5H-I**). CTCF^YF-AA^ depletion mainly downregulated genes that were strongly reduced by wild-type CTCF^AID^ depletion, while weakly downregulated genes were mostly unaffected (**Figure 3A**). For instance, of the genes downregulated more than twofold by CTCF^AID^ depletion, 85% (80 of 94) showed at least a 1.5-fold reduction in expression within 2 hours of CTCF^YF-AA^ depletion (**Figure 3B**). These commonly downregulated genes are mostly unaffected by A-485 treatment or RAD21^AID^ depletion. In sharp contrast, the genes (14/94) specifically downregulated by CTCF^AID^ but not by CTCF^YF-AA^ depletion are all downregulated by A-485 and tend to be downregulated after RAD21^AID^ depletion.

**Figure 3.**
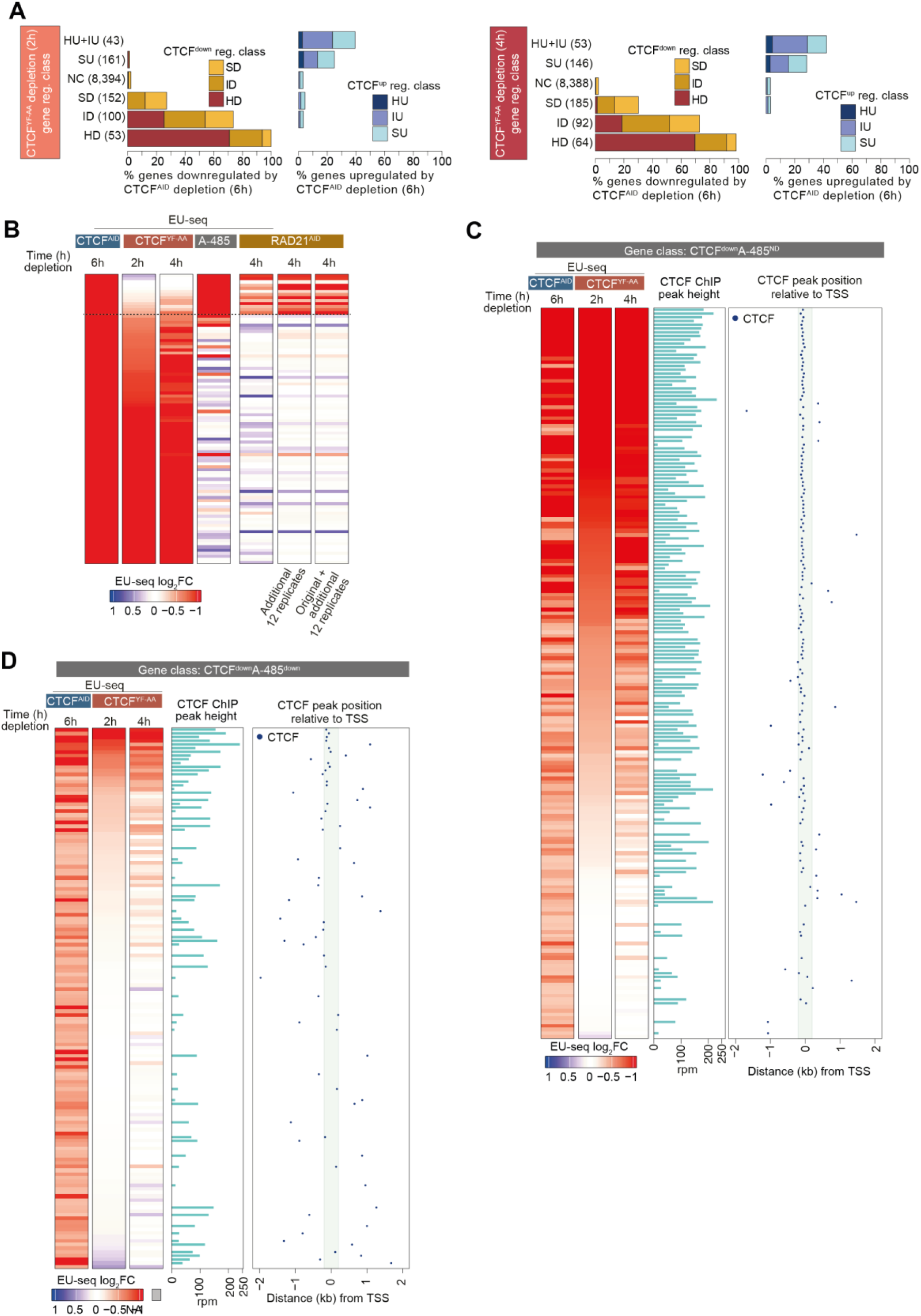
Promoter-bound CTCF activates genes independently of anchoring cohesin loops. **(A)** Cohesin interaction-defective CTCF^YF-AA^ can activate many CTCF-regulated genes. Genes regulated after acute (2h, left panels; 4h, right panels) CTCF^YF-AA^ depletion are grouped into indicated classes. Within each class, the fraction of genes regulated by the depletion of wild-type CTCF^AID^ is shown. CTCF^AID^ regulated gene class (CTCF^down^ reg. class, and CTCF^up^ reg. class) indicates the magnitude of gene regulation after depletion (6h) of CTCF^AID^. **(B)** CTCF^YF-AA^ lacks the ability to activate genes by cohesin-dependent looping. The heatmap shows the gene regulation by treatment with A-485 or depletion of CTCF^AID^, CTCF^YF-AA^, or RAD21^AID^. This analysis only includes strongly downregulated genes after CTCF^AID^ depletion (≥ 2-fold down, EU-seq expression TPM >15 in CTCF^AID^ cells). The dotted line indicates genes that are downregulated <1.5-fold after CTCF^YF-AA^ depletion. **(C-D)** CTCF^YF-AA^ activated genes are enriched for CTCF binding in their promoters. Genes downregulated after CTCF^AID^ depletion are grouped into: CTCF^down^A-485^ND^ (C) and CTCF^down^A-485^Down^ (D). Within each group, shown is the fold-change in gene regulation after depletion of CTCF^AID^ (6h) and CTCF^YF-AA^ (2h, 4h). Genes are sorted by their downregulation after CTCF^YF-AA^ depletion (2h), and heatmaps show the magnitude of downregulation in the indicated conditions. For the genes with CTCF binding +/- 2kb from TSS, the position of CTCF binding and height of CTCF ChIP peak are displayed. Region +/- 200bp from TSS is highlighted. See also Supplemental Figure 4.

Prompted by this result, we grouped CTCF^down^ genes based on their regulation by A-485 and examined their regulation upon CTCF^YF-AA^ depletion. In the CTCF^down^A-485^ND^ group, most genes were downregulated within 2 hours of CTCF^YF-AA^ depletion (**Figure 3C**). In contrast, in the CTCF^down^A-485^down^ group, CTCF^YF-AA^ depletion affected only a few genes (**Figure 3D**). Notably, in both groups, genes downregulated by CTCF^YF-AA^ were predominantly bound by CTCF in their promoters. These results demonstrate that: (1) CTCF^YF-AA^ retains activation ability for genes with promoter-bound CTCF but loses its capacity to drive activation through cohesin and CBP/p300-dependent enhancers. (2) Strongly downregulated genes in CTCF^YF-AA^-depleted cells do not require cohesin or CBP/p300 enhancers. (3) A small subset of CTCF^down^ genes may depend on both cohesin-dependent and independent CTCF functions.

**Supplemental Figure 5.**
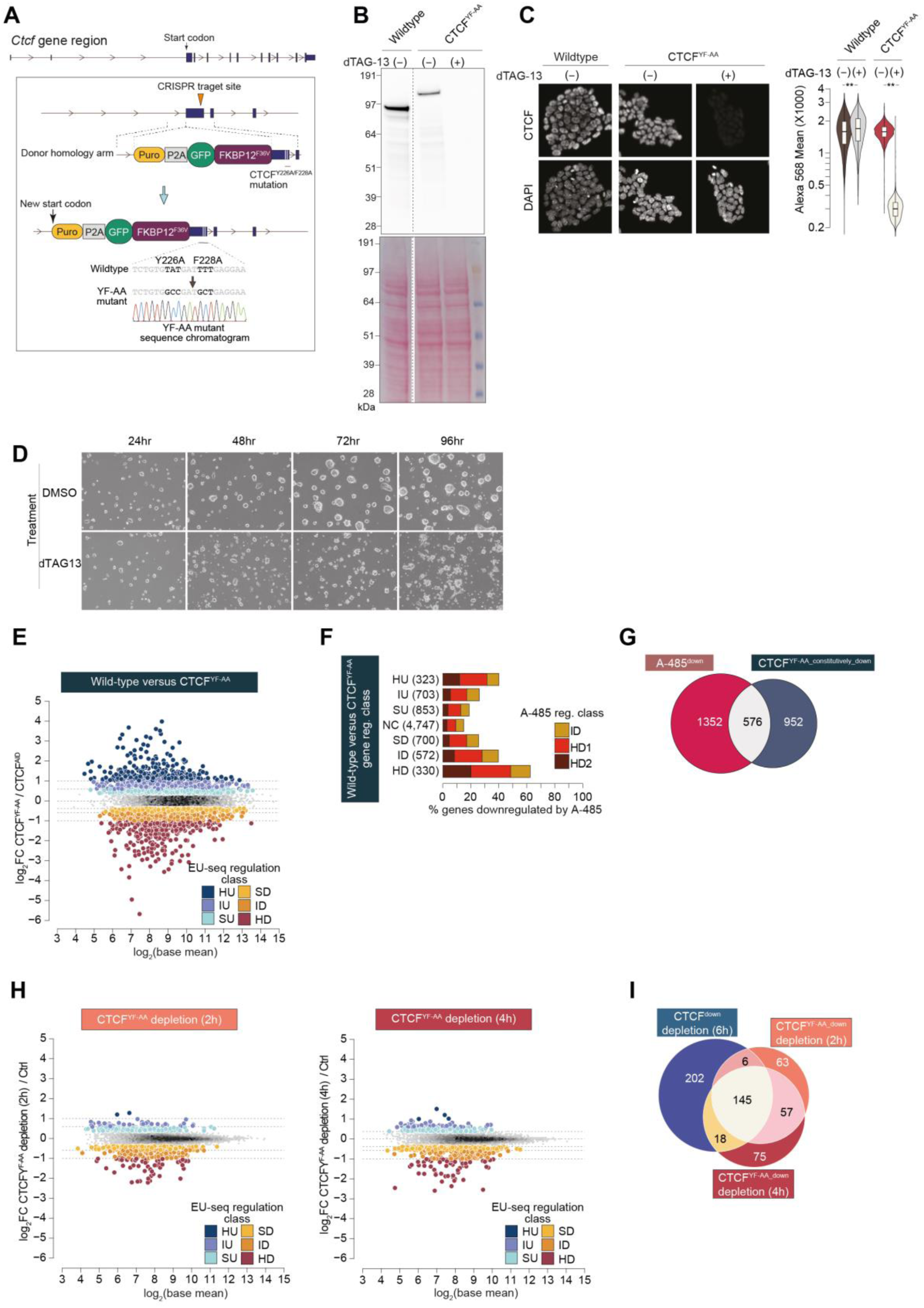
Acute depletion of CTCF^YF-AA^ impairs cell proliferation and downregulates a large portion of genes downregulated by CTCF^AID^ depletion, related to Figure 4. **(A)** Schematic for generating CTCF^YF-AA^ cells. Endogenous *Ctcf* is modified by fusing it with Puro-P2A-Gfp-Fkbp12^F36V^ and introducing Y226A and F228A mutations. Both mutations are confirmed by sequencing of the modified genomic region. **(B-C)** Confirmation of dTAG-13-induced depletion of CTCF^YF-AA^ by immunoblotting (B) and immunofluorescence (C). Ponceau staining serves as a protein loading control. CTCF expression in immunofluorescence is quantified by anti-CTCF antibody, and nuclei are stained using DAPI. The box plots display the mean CTCF fluorescence intensity distribution in each condition. The median, upper and lower quartiles; the whiskers show the 1.5× interquartile range (IQR) are indicated. Two-sided Mann–Whitney U-test, followed by correction for multiple comparisons with the Benjamini–Hochberg method; **Padj<0.001. **(D)** Micrographs showing colony growth of CTCF^YF-AA^ mESCs, with or without depletion of CTCF^YF-AA.^ Cells were imaged at the indicated time points after starting the dTAG-13 or DMSO treatments. **(E)** Gene expression changes in CTCF^YF-AA^ cells. Gene expression is compared between CTCF^YF-AA^ and CTCF^AID^ cells cultured under standard conditions without any treatments. **(F)** Differentially expressed genes in CTCF^YF-AA^ cells are classified into the indicated class. HU: >2-fold upregulated, IU: 1.5-2-fold upregulated, SU: 1.3-1.5-fold upregulated, NC: <1.2-fold change, SD: 1.3-1.5-fold downregulated, ID: 1.5-2-fold downregulated, HD: >2-fold downregulated. **(G)** Overlap between A-485 downregulated genes and genes downregulated in constitutively CTCF^YF-AA^ expressing cells. **(H)** Change in gene expression after depletion of CTCF^YF-AA^ for 2h (left panel) or 4h (right panel). Based on fold-change in gene expression, regulated genes are grouped into the indicated categories, as specified in Supplemental Figure 1B. **(I)** Overlap between genes downregulated after acute the depletion of CTCF^AID^ (6h), and after CTCF^YF-AA^ depletion (2h, or 4h).

### CTCF may function as a repressor, but seldomly

An early study using plasmid-based reporters suggested that promoter-bound CTCF acts as a transcriptional repressor ^49^. To investigate the in vivo relevance of this observation, we analyzed genes commonly upregulated by CTCF^AID^ and CTCF^YF-AA^ depletion and those upregulated solely by CTCF^AID^ depletion. Genes upregulated only by CTCF^AID^ depletion show nearby enhancer enrichment and tend to be downregulated by A-485 (**Supplemental Figure 6A**). In contrast, genes commonly upregulated by CTCF^AID^ and CTCF^YF-AA^ are more likely to have CTCF bound at or just downstream of the TSS (−50 to +400 bp) than genes upregulated only by CTCF^AID^ (31.4% vs. 6.7%) (**Supplemental Figure 6B-C**). These commonly upregulated genes lack strong proximal enhancers and are not notably downregulated by A-485. Together, our data supports in vitro data that CTCF binding in promoters may repress transcription^49,50^, but this appears to affect very few genes, suggesting that this is not a major function of CTCF.

### Promoter-bound CTCF activates housekeeping genes that are essential for cell proliferation

Next, we investigated the properties of genes regulated by promoter-bound CTCF and the conservation of promoter CTCF binding. RAD21^down^, A-485^down^, and CTCF^down^A-485^down^ genes show expression in restricted tissues (**Figure 4A**). In contrast, genes downregulated by both CTCF^YF-AA^ and CTCF^AID^ depletion but not by A-485 (CTCF^down^CTCF^YF-AA_down^A-485^ND^) are expressed across a broad range of tissues, with more than half displaying ubiquitous expression across 151 mouse tissues. Analysis of CTCF ChIP-seq data in diverse mouse and human tissues revealed that CTCF binding is significantly more conserved in promoters of CTCF^down^CTCF^YF-AA_down^A-485^ND^ genes than promoters of CTCF^NC^ and CTCF^down^A-485^down^ genes (**Supplemental Figure 6D**). This suggests a conserved role of promoter-bound CTCF in gene activation.

**Figure 4.**
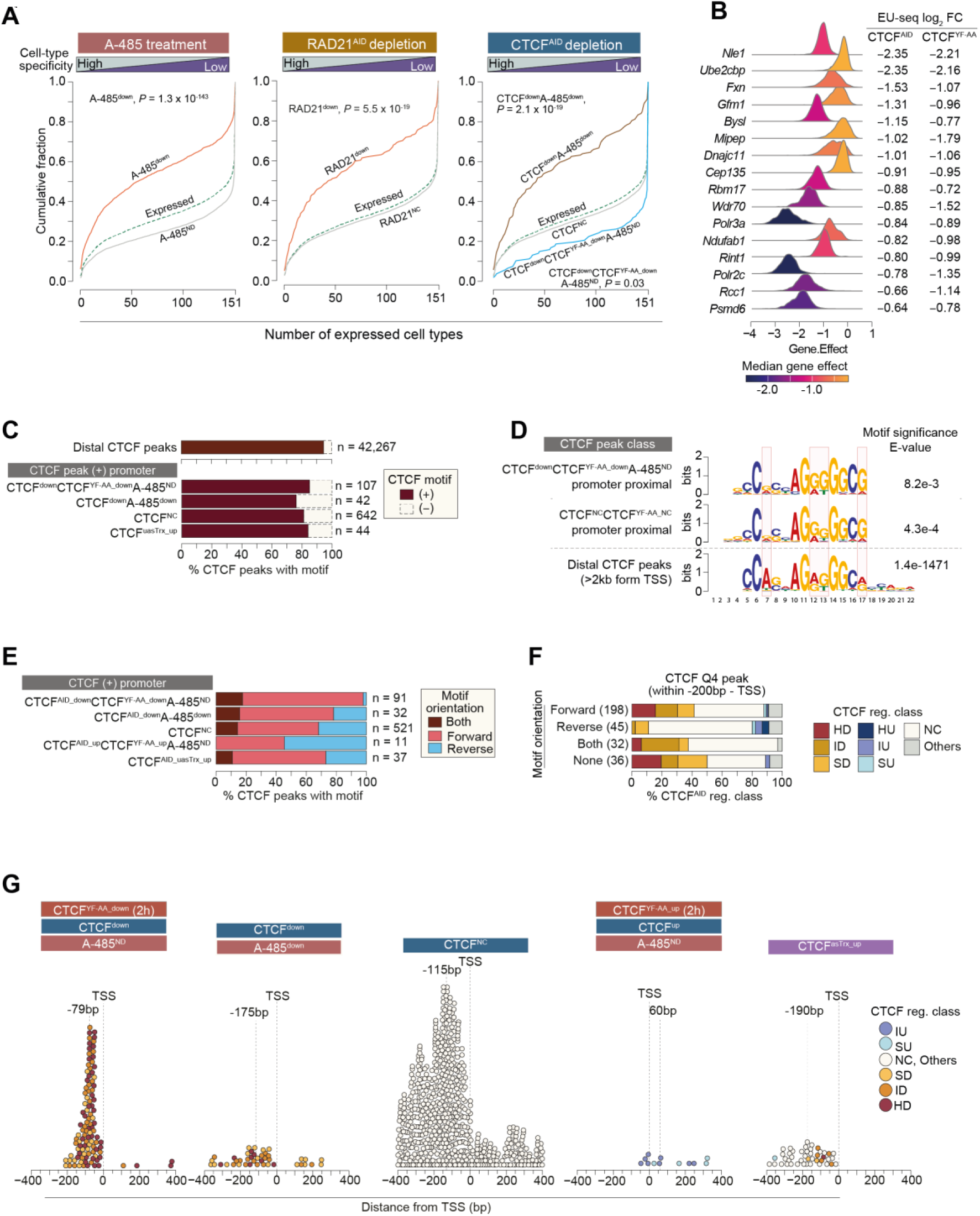
CTCF functions as a transcription activator for housekeeping genes, and its functional roles are encoded by positional binding. **(A)** Cell type specificity of the indicated classes of regulated genes. For the indicated groups, the expression of genes is analyzed across 151 mouse tissues and developmental stages. “Expressed” includes all expressed genes in mESC detected in the indicated conditions. P-values were calculated using a two-sample Kolmogorov-Smirnov test. **(B)** A list of common essential genes that are downregulated by both CTCF^AID^ and CTCF^YY-AA^ depletion but not by A-485. Common essential genes are defined by the DepMap database based on their essentiality for the proliferation of human cancer cell lines. **(C)** Fraction of CTCF peaks with detectable canonical CTCF binding motif. In the indicated classes of genes, CTCF motifs are searched in +/- 100bp from CTCF peak summit (peak summit distance: +/- 400bp from TSS). Distal (>2kb away from TSS) CTCF peaks are included as a reference. **(D)** Sequence motif enriched in CTCF peaks detected in distal and promoter (upstream 200bp – downstream 50bp from TSS) regions. **(E)** CTCF motif orientation within CTCF peak regions in the promoters (+/- 400bp from TSS) of the indicated class of regulated genes. **(F)** CTCF^AID^ depletion-induced regulation of genes strongly bound by CTCF (Q4 of CTCF peaks) in their promoters (upstream 200bp – TSS). Genes bound by Q4 CTCF peaks are grouped by CTCF motif orientation, and within each group, the fraction of the CTCF-regulated gene class is depicted. **(G)** Distribution of CTCF binding position in the promoter-proximal regions (+/- 400bp of TSS) of the indicated classes of genes. In each group, the dotted lines indicate the TSS position and the median CTCF peak distance from TSS. See also Supplemental Figures 5 and 6.

To investigate if genes activated by CTCF are required for cell proliferation, we checked for genes that are defined as common essential in the DepMap database (https://depmap.org/portal). CTCF^YF-AA^ and CTCF^AID^ depletion downregulate many ubiquitously essential genes, including *Nle1*, *Bysl*, *Rbm17*, *Wdr70*, *Polr3a*, *Polr2c*, and *Psmd6* (Figure 4B). This suggests that CTCF’s essentiality for mammalian cells is likely related to its role in activating essential housekeeping genes.

### Different CTCF functional outcomes are associated with distinct positional binding

Our findings indicate that promoter-bound CTCF plays a role in gene activation. However, promoter-bound CTCF has also been shown to repress upstream antisense transcription (uasTrx) without affecting the transcription of the gene’s sense strand ^51^. We examined CTCF-dependent uasTrx regulation in our data to clarify these paradoxical observations. Most upstream anti-sense transcripts are short and rapidly degraded, making their detection challenging in EU-seq data. Still, the computational analysis identified increased uasTrx expression (>2-fold; termed CTCF^uasTrx_up^) in 80 promoters following CTCF^AID^ depletion. In 60% of these promoters, CTCF binding and CTCF-dependent uasTrx expression were confirmed through manual data inspection and used for further analysis.

First, we assessed CTCF motif orientation in promoters. Notably, 18.6% of promoter-bound CTCF peaks lack canonical motifs, compared to only 5.3% in distal regions (**Figure 4C**), even though CTCF enrichment at promoters is comparable to or higher than at distal sites (**Supplemental Figure 4A-B**). De novo motif analyses reveal that CTCF motifs in promoters differ from non-promoter sites, particularly at positions 7, 12, 13, and 17 (**Figure 4D**). However, among promoter-bound CTCF sites, these motif differences were consistent regardless of gene downregulation after CTCF depletion. Instead, regulated gene groups exhibited notable differences in motif orientation: for example, only 2% of promoters in CTCF^down^CTCF^YF-AA_down^A-485^ND^ genes had exclusively reverse motifs, compared to 22% of CTCF^down^A-485^ND^, 27% of CTCF^uasTrx_up^, and 32% of CTCF^NC^ genes (**Figure 4E**).

To test whether CTCF motif orientation in promoters predicts gene regulation, we analyzed genes with strong CTCF binding near the TSS (+/- 200 bp). Among genes with forward-oriented or absent CTCF motifs, 29% were downregulated (>1.5-fold) after CTCF^AID^ depletion, while only 2% of genes with exclusively reverse motifs showed similar downregulation (**Figure 4F**). Promoters with reverse motif orientation exhibit no specific bias for downregulation; instead, they show a weak, similar tendency of up- and down-regulation after CTCF depletion.

The positioning of CTCF binding within promoters also correlated with functional outcomes (**Figure 4G**). For CTCF^down^CTCF^YF-AA_down^A-485^ND^ genes, CTCF binding occurs almost exclusively within 200 bp upstream of the TSS, with a median distance of approximately 80 bp. By contrast, in CTCF^down^A-485^down^, CTCF^uasTrx_up^, and CTCF^NC^ genes, CTCF binding was more dispersed and situated further upstream. Only a few genes show uasTrx upregulation and concomitant downregulation of the corresponding sense transcript, and they have CTCF binding closer to the TSS than in genes exhibiting only uasTrx upregulation (**Figure 4G**).

Together, the results confirm forward-oriented CTCF binding in CTCF^down^ gene promoters ^11,22^. Importantly, our results reveal that this motif orientation bias is extreme for CTCF^down^A-485^ND^ genes, which are mainly activated independently of cohesin and CBP/p300-dependent enhancers.

**Supplemental Figure 6.**
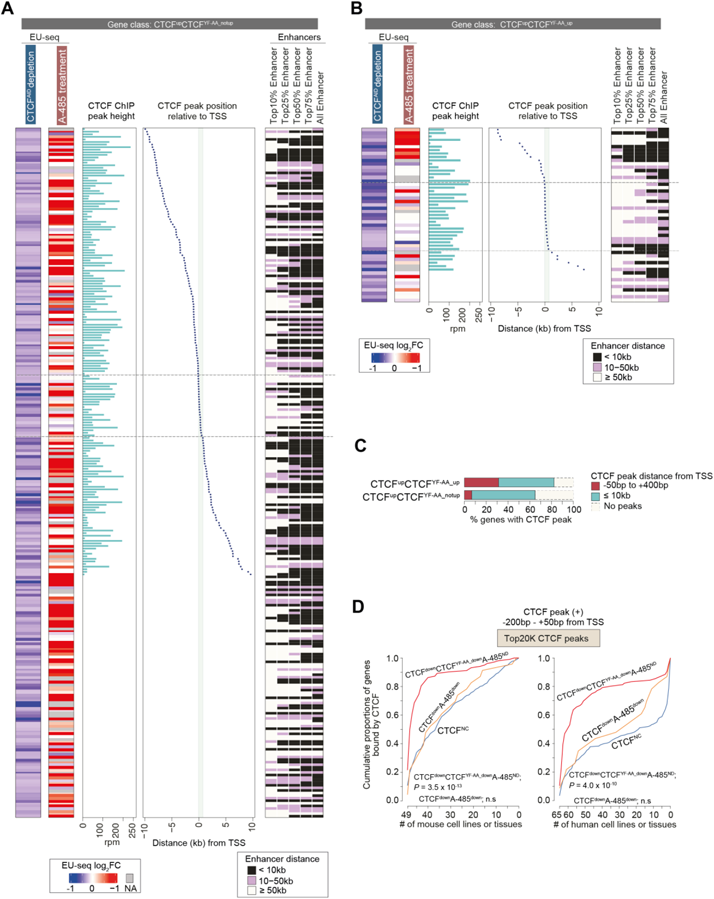
CTCF^YF-AA^ depletion increases expression of only a subset of CTCF^up^ genes and genes downregulated by CTCF^AID^ depletion show conserved CTCF binding in their promoters across diverse cell types, related to Figure 4. **(A)** Properties of genes upregulated (>1.3 fold) after acute (6h) CTCF^AID^ depletion, but not upregulated after acute (2h) CTCF^YF-AA^ depletion (CTCF^up^CTCF^YF-AA_notup^). The fold increase in gene expression after CTCF^AID^ depletion and the regulation of the same genes by A-485 treatment. CTCF peak position (+/- 2kb from TSS) and peak height are indicated, along with the enrichment of enhancers in their proximity. Enhancers are defined as overlapping regions of H2BK20ac peaks and H3K27ac peaks, excluding regions +/- 500bp from TSS. Enhancers are ranked by H2BK20ac ChIP signal enrichment. Regions +/- 200bp from TSS are highlighted. **(B)** Properties of genes commonly upregulated (>1.3-fold) after acute (6h) CTCF^AID^ depletion and after acute (2h) CTCF^YF-AA^ depletion (CTCF^up^CTCF^YF-AA_up^). Shown is the fold increase in gene expression after CTCF^AID^ depletion and regulation of the same genes by A-485 treatment. CTCF peak position (+/- 2kb from TSS) and peak height are indicated, along with the enrichment of enhancers in their proximity. Enhancers are defined as overlapping regions of H2BK20ac peaks and H3K27ac peaks, excluding regions +/- 500bp from TSS. Enhancers are ranked by H2BK20ac ChIP signal enrichment. Regions +/- 200bp from TSS are highlighted. **(C)** Fraction of genes showing CTCF binding within the indicated distance from TSS, for the indicated groups of regulated genes. Of note, genes commonly upregulated by CTCF^AID^ and CTCF^YF-AA^ depletion exhibit a higher prevalence of CTCF binding in promoter overlapping and immediate downstream regions. **(D)** Conservation of CTCF binding (upstream 200bp – downstream 50bp from TSS) in the promoters of indicated classes of CTCF-regulated genes in mouse (left panel) and human (right panel) cell lines and tissues. Only genes bound by CTCF in promoters of mESC are included in this analysis. P-values were calculated using a two-sample Kolmogorov-Smirnov test.

### Cohesin-loop anchoring defective CTCF acts as a canonical transcription activator

To investigate whether CTCF plays a direct role in promoter activation, we used a plasmid-based luciferase reporter assay to examine the promoters of eight CTCF^down^ genes bound by CTCF in their promoters (*Mrps2*, *Psmd6*, *Rabggta*, *Dusp12*, *Mrpl23*, *Polr2c*, *Tab1*, *Ttc5*) (**Supplemental Figure 7A**). Promoter activity was examined for wild-type sequences and mutant promoters lacking the putative CTCF binding site or having the CTCF motif sequence inverted. Also, we analyzed promoters of two CTCF^up^ genes (*Lsm11*, *Ocel1*), without or with mutation of CTCF binding motif. As controls, we included two promoters (*Psmd4* and *Actn4*) lacking CTCF binding and not regulated upon CTCF depletion. To further ensure that the measured reporter activity was attributable to the cohesin-independent function of CTCF, promoter activity was assessed in mESCs expressing CTCF^YF-AA^, both with and without acute (6h) depletion of CTCF^YF-AA^.

Notably, the activity of all eight CTCF^down^ gene promoters decreased significantly after CTCF^YF-AA^ depletion (**Figure 5A**). In six promoters (*Mrps2*, *Psmd6*, *Rabggta*, *Mrpl23*, *Polr2c*, *Ttc5*), mutation of the putative CTCF binding motif reduced activity. CTCF motif mutant *Dusp12* and *Tab1* promoters, and CTCF motif inverted *Tab1* promoter, remained dependent on CTCF^YF-AA^ expression. Both these promoters lacked canonical CTCF binding motifs, and the mutations were introduced based on manual motif inference. The sustained dependency of these promoters on CTCF suggests that CTCF may bind to an additional and/or different sequence than the one mutated. Regardless, the results show that CTCF^YF-AA^ can activate promoters of all (8/8) CTCF^down^ genes analyzed.

**Figure 5.**
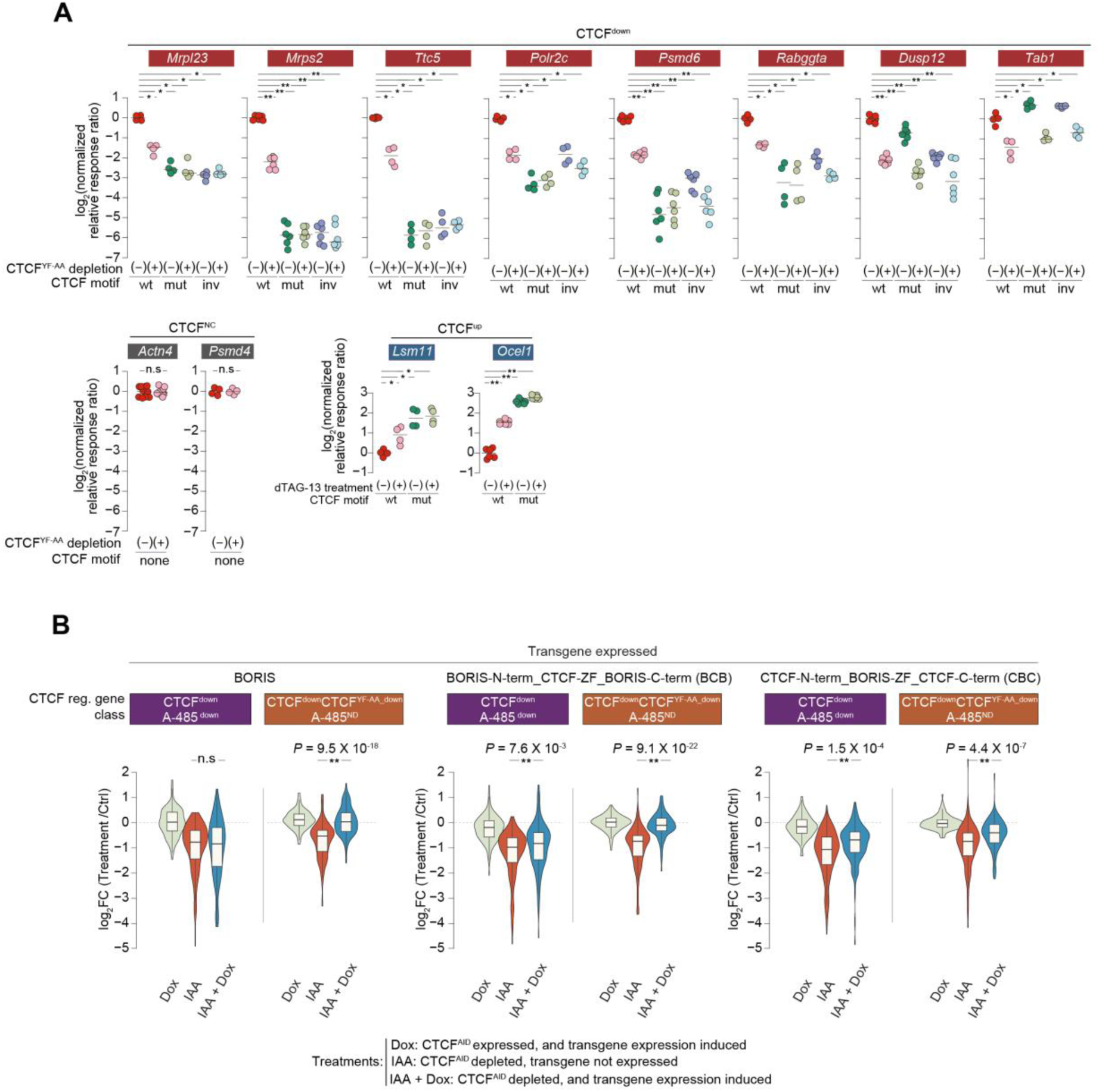
CTCF directly activates promoters, and CTCFL can activate a subset of CTCF targets. **(A)** Shown is the luciferase relative response ratio of the indicated CTCF^down^ gene promoters. Promoter activity is measured for wildtype (wt) sequences, putative CTCF binding mutant (mut) sequences, and putative CTCF binding motif inverted (inv) sequences. The reporter activity is measured in mESCs expressing ^dTAG^CTCF, with (+) or without (-) depleting CTCF^YF-AA^ by treating with dTAG-13 (6h). In each group, a horizontal line indicates the median value. Two-sided Mann–Whitney U-test, followed by correction for multiple comparisons with the Benjamini–Hochberg method; **Padj<0.001, *Padj<0.05. **(B)** Regulation of the indicated classes of CTCF downregulated genes after the overexpression of the indicated transgenes (Doxycyclin, Dox) in CTCF^AID^ cells, after depletion (24h) of CTCF^AID^ by IAA (indole-3-acetic acid), and after the depletion of CTCF^AID^ and simultaneous overexpression of the transgenes (IAA + Dox). The box plots display the median, upper and lower quartiles; the whiskers show the 1.5× interquartile range (IQR). Two-sided Mann–Whitney U-test, followed by correction for multiple comparisons with the Benjamini–Hochberg method; **Padj<0.001, *Padj<0.05. See also Supplemental Figure 7.

The activity of CTCF-independent gene promoters (*Psmd4*, *Actn4*) remained unaffected after CTCF^YF-AA^ depletion (**Figure 5A**). The CTCF^up^ gene promoters (*Lsm11*, *Ocel1*) showed increased activity after CTCF^YF-AA^ depletion. The basal activity of the mutant *Lsm11* and *Ocel1* promoters was higher than their wild-type counterparts, and their activity remained unchanged after CTCF^YF-AA^ depletion.

These results demonstrate a direct role of CTCF as a transcription activator and imply a role of motif orientation for promoter activation.

### The transcription activator function is interchangeable between CTCF and BORIS

CTCF has a vertebrate-specific, called BORIS (CTCFL), is usually only expressed in the testis, but aberrant expression is found in cancers ^52^. CTCF and BORIS share high sequence identity in their CTCF in DNA binding zinc fingers (ZFs) but exhibit minimal conservation in non-DNA binding N- and C-termini ^53^.

BORIS cannot anchor cohesin loops, but it can bind to thousands of sites occupied by CTCF ^54^. BORIS preferentially binds in promoters compared to distal CTCF sites ^55,56^, prompting us to investigate whether BORIS can regulate the transcription of CTCF-activated genes.

Deploying the same CTCF^AID^ in mESCs used here, a previous study conditionally swapped CTCF^AID^ with BORIS or chimeric constructs: CTCF-N-terminus_BORIS-ZF_CTCF-C-terminus (CBC) or BORIS-N-terminus_CTCF-ZF_BORIS-C-terminus (BCB) ^42^. Although different transgenes caused varied gene expression patterns, none fully restored genes dysregulated by CTCF^AID^ depletion.

Having identified enhancer-dependent and -independent functions of CTCF, we re-analyzed their data to explore how BORIS impacts their expression specifically. We categorized CTCF^down^A-485^down^ genes as likely enhancer-dependent CTCF targets and CTCF^down^CTCF^YF-AA_down^A-485^ND^ genes as likely enhancer-independent targets. Remarkably, in CTCF^AID^-depleted cells, overexpression of BORIS robustly and selectively restored expression of CTCF^down^CTCF^YF-AA_down^A-485^ND^ genes, but not CTCF^down^A-485^down^ genes (**Figure 5B**). The BCB transgene also selectively restored CTCF^down^CTCF^YF-AA_down^A-485^ND^ genes, albeit to a slightly lesser extent than BORIS, likely due to the weaker induction of the BCB transgene compared to BORIS (**Supplemental Figure 7C**). The expression of CBC partially restored the expression of both CTCF^down^CTCF^YF-AA_down^A-485^ND^ and CTCF^down^A-485^down^ genes. The partial restoration by CBC may reflect the differences in genomic occupancy of BORIS and CTCF ^55,56^ and only partial (∼50%) restoration of cohesin loops by chimeric CTCF-N-terminus-BORIS ^54^. Regardless, these findings show that BORIS can selectively restore the expression of enhancer-independent CTCF targets, demonstrating that both BORIS and CTCF are competent transcription activators. This finding may help explain why BORIS preferentially binds in gene promoters compared to distal regions ^55,56^.

**Supplemental Figure 7.**
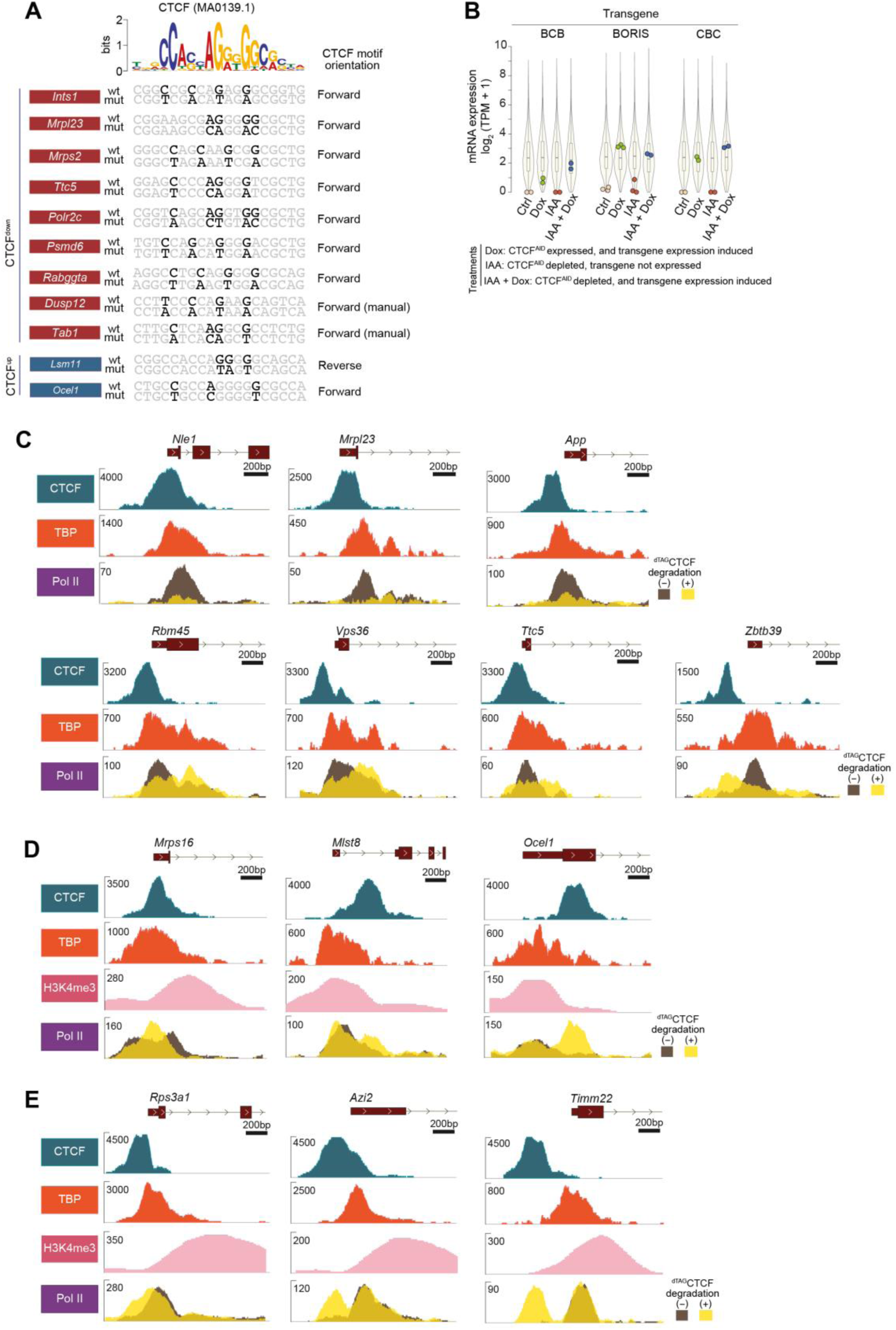
Functional and mechanistic consequence of promoter CTCF binding for gene regulation, related to Figures 5 and 6. **(A)** Displayed are sequences of putative CTCF binding motifs within the promoters of specified CTCF-regulated genes. The motif at the top represents the canonical CTCF binding motif. Nucleotides in dark black type denote mutations aimed at disrupting CTCF binding in the putative motifs. CTCF motif orientation is with respect to the orientation of the indicated genes in the genome. For *Dusp12* and *Tab1* promoters, computational analyses did not identify canonical motifs within CTCF peak regions; hence, the provided sequences represent manually inferred putative CTCF binding sequences. **(B)** Luciferase relative response ratio of the indicated CTCF^NC^ (*Actn4, Psmd4*) gene promoters. Promoter activity is measured in mESCs expressing ^dTAG^CTCF, with (+) or without (-) depleting CTCF^YF-AA^ by treating with dTAG-13 (6h). In each group, the horizontal line indicates the median value. Two-sided Mann–Whitney U-test, followed by correction for multiple comparisons with the Benjamini–Hochberg method; **Padj<0.001, *Padj<0.05, n.s., not significant. **(C)** Expression of the specified transgenes under designated treatment conditions: Doxycycline (Dox) treatment induces transgene expression, while indole-3-acetic acid (IAA) treatment depletes endogenous CTCF^AID^ protein. All transgenes are mRuby2-fused, ensuring expression quantification via sequence read counts mapping to mRuby2, avoiding confounding by endogenous gene transcripts (e.g., for *Ctcf*). TPM distribution of all the expressed genes is shown as a violin plot. The box plots display the median, upper and lower quartiles; the whiskers show the 1.5× interquartile range (IQR). CBC: CTCF-N-terminus_BORIS-ZF_CTCF-C-terminus, BCB: BORIS-N-terminus_CTCF-ZF_BORIS-C-terminus. Gene expression data sourced from reference ^42^. **(D-F)** Shown are genome browser tracks of representative CTCF^down^ genes (**D**), CTCF^up^ genes (**E**), and CTCF^uasTrx_up^ genes (**F**) after acute (6h) depletion of CTCF. Pol II enrichment is analyzed with or without depletion (4h) of ^dTAG^CTCF. ChIP-seq profiles of CTCF, TBP, and where specified H3K4me3, are included as references. Among the shown genes, *App* and *Ocel1* are expressed at a low level (EU-seq TPM <15).

### CTCF controls chromatin accessibility

To further understand how CTCF exerts distinct functions, we analyzed CTCF’s role in chromatin accessibility. To demonstrate the robustness of gene expression changes detected in CTCF^AID^ cells, we generated an independent CTCF depletion cell line (*Gfp-Fkbp12^F36V^-Ctcf*; hereafter ^dTAG^CTCF). Change in chromatin accessibility quantified after acute (3h) depletion of ^dTAG^CTCF depletion. Globally, chromatin accessibility was reduced (>2-fold) in 6.4% of regions but increased at only 0.03% (**Figure 6A**). This contrasts sharply with the widespread, equally prominent gain and loss in chromatin accessibility after long-term (24h) CTCF^AID^ depletion ^57^.

**Figure 6.**
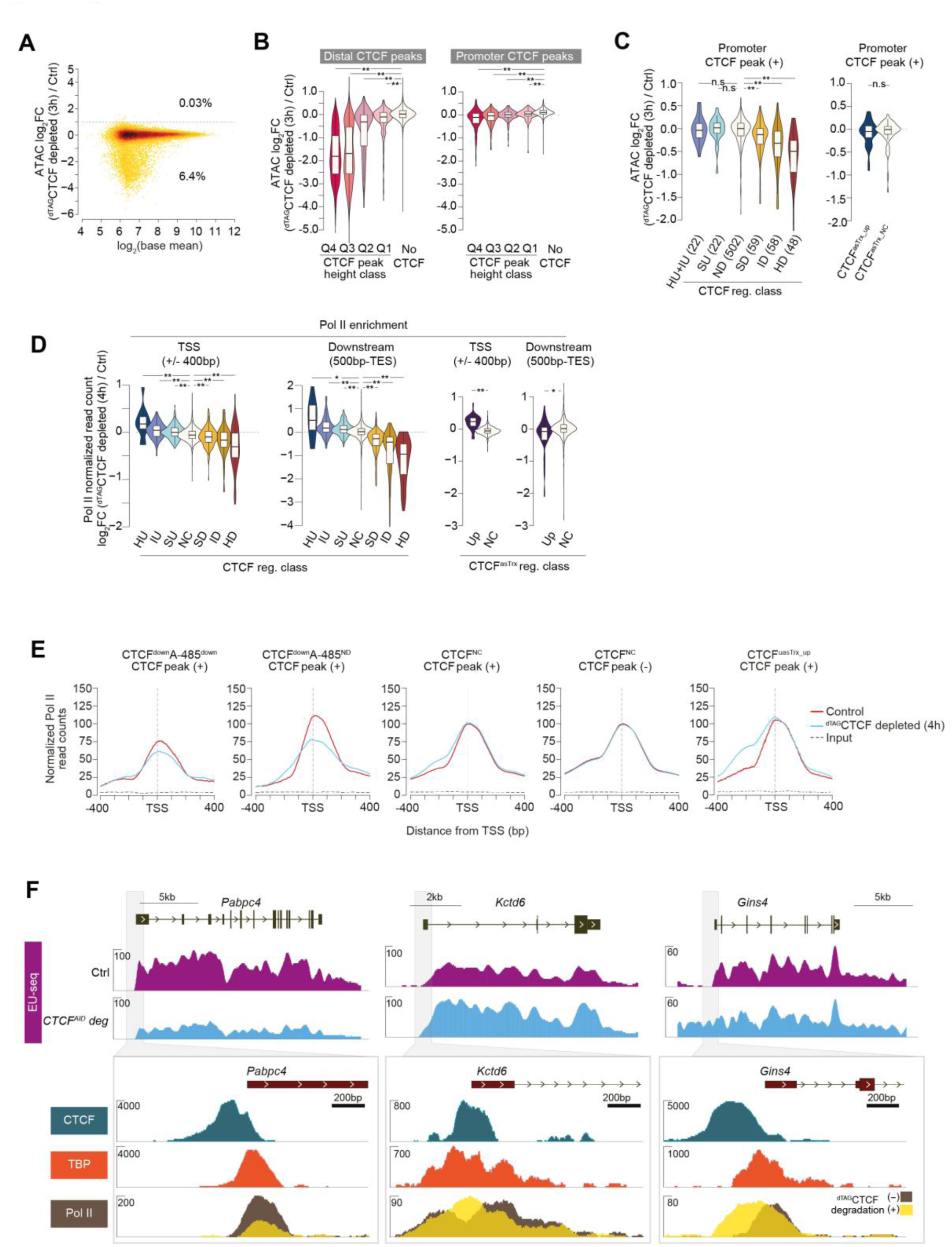
CTCF binding in promoters regulates chromatin accessibility and the recruitment and positioning of Pol II. **(A)** Change in global chromatin accessibility after ^dTAG^CTCF depletion. The percentage of ATAC-seq peaks showing a ≥2-fold change in accessibility is indicated. Change in chromatin accessibility is quantified 3h after ^dTAG^CTCF depletion. **(B)** Change in chromatin accessibility in CTCF peak regions after ^dTAG^CTCF depletion. Distal (>200bp from TSS) and promoter (+/- 200bp from TSS) ATAC peaks are grouped into quartiles (Q1-Q4 and No) based on overlapping CTCF peak height. Two-sided Mann–Whitney U-test, followed by correction for multiple comparisons with the Benjamini–Hochberg method; **Padj<0.001, *Padj<0.05. **(C)** In the indicated CTCF-regulated gene class, the change in chromatin accessibility after ^dTAG^CTCF depletion. The analysis only includes promoters bound by CTCF (within +/-200bp of TSS). In the left and right panels, promoters are classified based on the regulation of sense or anti-sense transcript, respectively, after ^dTAG^CTCF depletion. Two-sided Mann–Whitney U-test, followed by correction for multiple comparisons with the Benjamini–Hochberg method; **Padj<0.001, *Padj<0.05. **(D)** Change in Pol II binding in the indicated groups of CTCF-regulated gene class. Pol II enrichment is quantified in the promoter (+/-400bp from TSS) and non-promoter (500bp-transcript end site (TES) regions. Normalization was performed based on the assumption that the Pol II binding at the gene body region of the CTCF non-regulated genes does not change between the control and ^dTAG^CTCF depleted condition. Change in Pol II binding is analyzed by ChIP-seq after ^dTAG^CTCF depletion (4h). The box plots display the median, upper and lower quartiles; the whiskers show the 1.5× interquartile range (IQR). Two-sided Mann–Whitney U-test, followed by correction for multiple comparisons with the Benjamini–Hochberg method; **Padj<0.001, *Padj<0.05. **(E)** Pol II binding in promoters of the indicated groups of CTCF-regulated genes, with (+) or without (-) CTCF binding in their promoters. **(F)** Upper genome browser tracks show representative genes exhibiting decreased (*Pabpc4*) or increased (*Kctd6*) transcription or increased upstream anti-sense transcription (*Gins4*). Lower panels show the binding of CTCF, TBP, and Pol II in the promoters of indicated genes. Pol II binding is analyzed with or without depletion of ^dTAG^CTCF. For additional examples see Supplemental Figures 7D-F. See also Supplemental Figure 7.

Overall, accessibility was more strongly reduced at distal peaks than promoters, and the degree of reduction corresponded with the extent of CTCF enrichment (**Figure 6B**). In promoters, accessibility was specifically reduced at CTCF^down^ genes that were bound by CTCF, and the decrease in accessibility corresponded with the reduction in gene transcription (**Figure 6C**). Interestingly, accessibility was not reduced in CTCF^NC^ and CTCF^uasTrx_up^ genes, even though CTCF binds their promoters. Together, the results confirm CTCF’s role in maintaining nucleosome-depleted regions ^58^ and demonstrate that CTCF regulates accessibility in only a subset of promoters, and the genes where CTCF regulates accessibility are rapidly downregulated after CTCF depletion.

### CTCF is a position-specific transcription activator and repressor

Next, we investigated how CTCF depletion affects the recruitment of RNA polymerase II (Pol II). Upon ^dTAG^CTCF depletion, Pol II recruitment significantly decreases in CTCF^down^ genes, correlating with the extent of gene downregulation observed in EU-seq (**Figure 6D**). The reduction in Pol II binding appears more pronounced in TSS downstream regions compared to TSS proximal regions (+/- 400bp from TSS). Conversely, in CTCF^uasTrx_up^ genes, Pol II binding increases in TSS proximal areas but not in downstream gene regions (**Figure 6D**). CTCF also subtly affects the position of Pol II binding (**Figure 6E**). Depletion of ^dTAG^CTCF results in an upstream shift in Pol II binding in CTCF^uasTrx_up^ genes and in CTCF^NC^ genes, albeit to a lesser extent. This shift is directly attributable to CTCF, as genes lacking CTCF binding in their promoters show no change in Pol II position.

While CTCF’s role in Pol II recruitment is already apparent in global analyses, examining individual genes reveals intricate details (**Figure 6F**, **Supplemental Figure 7D-F**). In most CTCF^down^ genes, such as *Pabpc4*, *Nle1*, and *Mrpl23*, CTCF depletion decreases Pol II recruitment in promoters without a notable change in Pol II binding position. Reassuringly, Pol II binding is reduced in the promoter of *App*, the first promoter to be shown to be activated by CTCF in plasmid-based reporter assay ^59^, although this gene is expressed weakly (EU-seq TPM <15) in mESC. In a small subset of CTCF^down^ genes, including *Rbm45*, *Vps36*, *Ttc5*, and *Zbtb39*, Pol II recruitment is reduced weakly, and the binding appears to shift downstream or upstream of the TSS.

CTCF appears to repress transcription in a few genes, and they are upregulated after CTCF^AID^ and CTCF^YF-AA^ depletion (**Figure 6F**, **Supplemental Figure 7E**). In this group, some genes have CTCF bound within the core promoter (*Mrps16*, *Kctd6*), indicated by the overlap with TBP binding. Pol II binding is not increased noticeably in these genes, but the binding shifts towards the core promoter after ^dTAG^CTCF depletion. In other genes (*Ocel1*, *Mlst8*), CTCF binds immediately downstream of TBP. In these genes, ^dTAG^CTCF depletion leads to increased Pol II binding in the TSS downstream region. Intriguingly, *Ocel1* and *Mlst8* show an unusual H3K4me3 pattern. Usually, the H3K4me3 marks the +1 nucleosome, and the H3K4me3 peak lies downstream of TBP, but in these genes, H3K4me3 is shifted upstream and overlaps with the TBP peak.

Similar to CTCF-upregulated genes, in CTCF^uasTrx_up^ genes, ^dTAG^CTCF depletion results in increased Pol II binding (**Figure 6F**, **Supplemental Figure 7F**). In genes where CTCF binds immediately upstream to promoters (*Gins4*, *Rps3a1*), Pol II binding is shifted upstream, but the Pol II peak appears as one contiguous peak. However, if CTCF binds further upstream of the promoter (*Timm22*, *Azi2*), ^dTAG^CTCF depletion does not result in the broadening of the original promoter Pol II peak; instead, ^dTAG^CTCF depletion leads to the appearance of a new Pol II peak.

Notably, both in CTCF^uasTrx_up^ and CTCF^up^ genes, ^dTAG^CTCF depletion results in increased accumulation of Pol II binding precisely in the position occupied by CTCF (**Figure 6F, Supplemental Figure 7E-F**). The only difference is that in CTCF^uasTrx_up^ genes, CTCF binds upstream of TSS so that Pol II binding is increased in the TSS upstream region, whereas in CTCF^up^ genes, CTCF binds within or downstream of the core promoter, and Pol II binding is increased within promoter or downstream region. This indicates that CTCF binding possibly occludes Pol II binding or hinders elongation. Depending on the position of CTCF binding relative to the TSS, CTCF can act as a repressor of either anti-sense or sense transcript.

Together, these findings reveal that depending on its precise location in the promoter, CTCF serves two opposing roles: it either facilitates or hinders Pol II recruitment and/or elongation. This suggests that CTCF acts as a position-specific transcription activator or repressor.

### Cohesin promotes CBP/p300-dependent gene activation in differentiated cells

3D genome organization differs markedly between pluripotent and differentiated cells ^60^. To explore cohesin’s role in gene activation in differentiated cells, RAD21^AID^ mESCs were differentiated into neuronal progenitor cells (NPCs). Transcriptional changes were measured after A-485 treatment for 1 hour and RAD21^AID^ depletion for 1, 2, and 4 hours.

The number of downregulated genes increases with prolonged RAD21^AID^ depletion, and genes regulated at different time points show strong overlap (**Figure 7A**, **Supplemental Figure 8A-C**). Similar to mESCs, RAD21^down^ genes in NPCs are heavily biased towards A-485-downregulated genes, while RAD21^up^ genes show minimal bias (**Figure 7B**). Among A-485-downregulated genes, those with stronger A-485 downregulation are more likely to be affected by RAD21 depletion: 42-51% of genes showing >4-fold downregulation by A-485 also exhibited decreased expression within 2-4 hours of RAD21^AID^ depletion. In contrast, genes unaffected by A-485 were largely resistant to RAD21 depletion.

**Figure 7.**
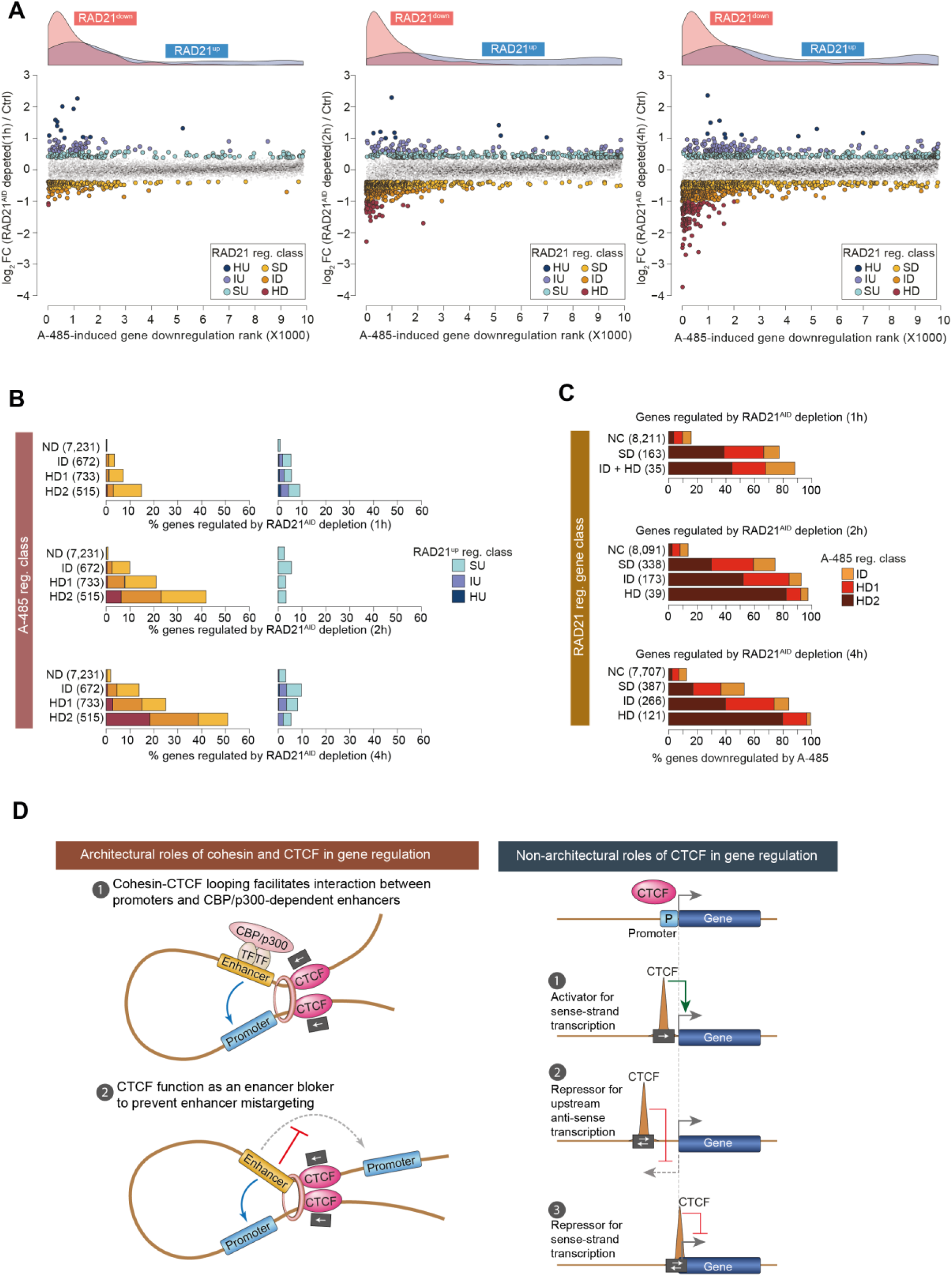
Position-specific binding of CTCF in promoters distinctly regulates Pol II recruitment and positioning. **(A)** Fold-change in gene expression in NPC after acute (1-4h) depletion of. Genes are ranked by A-485-induced fold downregulation, and the density plots (top) show the distribution of genes up- and down-regulated after RAD21^AID^ depletion. Genes up- and down-regulated after RAD21^AID^ depletion are classified as specified in Supplemental Figure 8A. **(B)** Within A-485 regulated gene groups, fraction of genes up- and down-regulated after RAD21^AID^ depletion in NPC. A-485 regulated genes are grouped into the indicated categories, within each category, the fraction of genes upregulated or downregulated after acute (1h, 2h, and 4h) depletion of RAD21^AID^ is depicted. **(C)** Within A-485 regulated gene groups, the fraction of genes downregulated by A-485 treatment in NPC. Genes regulated after RAD21^AID^ depletion are grouped into the indicated classes, and within each class, the fraction of genes regulated by A-485 is shown. **(D)** Proposed model for the architectural and non-architectural roles of cohesin and CTCF in gene regulation. In their architectural roles, cohesin and CTCF create chromatin loops that facilitate interactions between CBP/p300-dependent enhancers and their distal target promoters. CTCF also functions as an enhancer blocker, preventing mistargeting of enhancers to non-target promoters. Independently of its architectural role, CTCF regulates gene expression through position- and orientation-specific binding near TSS. CTCF acts as a transcription activator when bound immediately upstream of the TSS, represses upstream antisense transcription when positioned further upstream, and blocks transcription on the sense strand when bound within the core promoter or just downstream of it. The gene activator function of CTCF is strongly associated with its binding in the forward orientation, while motif orientation appears less critical for repressing sense and antisense transcription. See also Supplemental Figures 8 and 9.

Reciprocally, among the genes strongly (>2-fold) downregulated after 2-4 hours of RAD21^AID^ depletion, 92-97% are also downregulated by A-485 to the same extent (i.e. >2-fold down) (**Figure 7C**). Only 5% of genes in NPC are downregulated by A-485 by >4-fold (**Supplemental Figure 8B**), but among the genes strongly (>2-fold) downregulated after 2-4 hours of RAD21^AID^ depletion, 80-82% are downregulated by >4-fold by A-485 (odds ratio Inf, p < 2.2e-16, down-regulated versus ND, Fisher’s exact test). CTCF and RAD21 binding is enriched within 2 kb of RAD21^down^ genes (**Supplemental Figure 8D**).

CBP/p300-regulated genes in NPC include several known and putative enhancer targets, such as *Myc*, *Gli3*, *Kitl*, *Fbn2*, *Enc1*, and *Tnfrsf21*. Some of these genes, such as *Myc*, *Kitl*, *Enc1*, *Tnfrsf21*, and *Fbn2,* are downregulated both by RAD21^AID^ depletion and A-485 treatment, whereas others, such as *Gli2* and *Zfp608*, are only downregulated by A-485, not by RAD21^AID^ depletion (**Supplemental Figure 9A**).

Notably, A-485 treatment downregulates both *Gli3* and the proximal gene, *Inhba*, but RAD21^AID^ depletion only downregulates *Inhba*. A recent study identified a new type of scaffolding element in the *Zfp608* ^61^. which serves a dual function as an enhancer and architectural element to specifically promote *Zfp608* looping with multiple candidate enhancers in NPC but not in mESC ^61^. Surprisingly, deletion of this element disrupts enhancer-promoter interactions in NPC, but the expression *Zfp608* remains unchanged. In our data, *Zfp608* is downregulated by A-485, supporting the idea that it is activated by enhancers (**Supplemental Figure 9A**). However, *Zfp608* expression is not reduced after RAD21^AID^ depletion, showing that cohesin-dependent looping is dispensable for the activation of this gene, at least in the tested conditions and timeframe.

For some of the RAD21^down^ genes in NPC, cohesin-dependent loops are visible in Micro-C data from mESCs (**Supplemental Figure 9B**). Cohesin appears to promote looping in the *Tnfrsf21* and *Enc1*^20^ loci in mESC, and both of these genes are strongly regulated by RAD21^AID^ depletion in NPC. Also, the cohesin-dependent looping is observed in the *Fbn2* locus ^5,62^, even though this gene is not detectably expressed in mESC. *Fbn2* is robustly expressed in NPC and downregulated by both RAD21^AID^ depletion and A-485 treatment. These anecdotal examples support the role of looping in gene activation and indicate that cohesin-dependent domains may pre-exist before the commissioning of enhancers and their target genes.

Together, these results suggest that cohesin may play an even more prominent role in regulating genes in NPC than in mESC, and cohesińs preference for promoting activation of CBP/p300-dependent genes remains conserved between cell types.

**Supplemental Figure 8.**
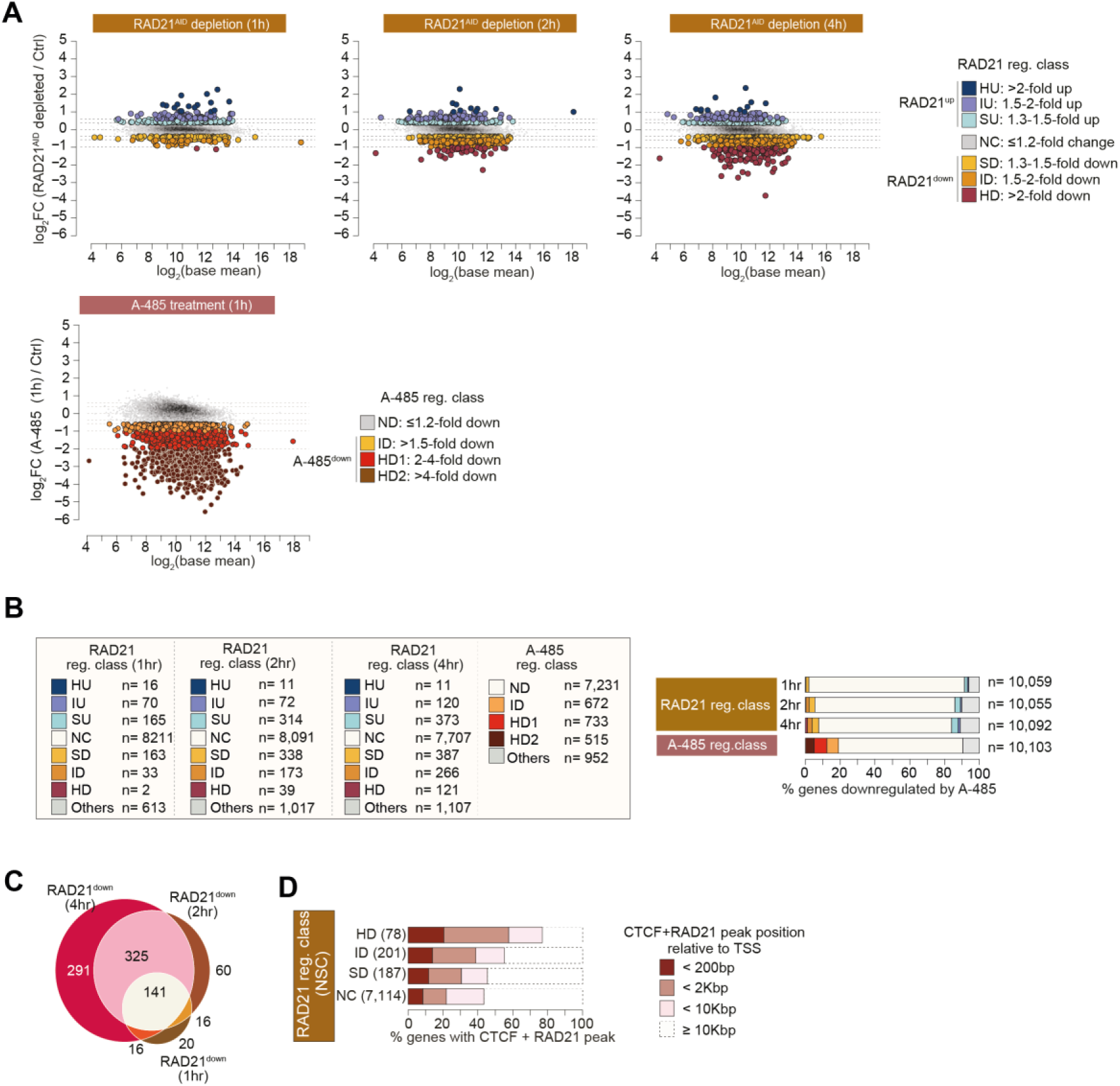
Acute RAD21^AID^ depletion downregulates hundreds of genes in NPC, related to Figure 7. **(A)** Transcription changes after RAD21^AID^ depletion in NPC. mESC expressing RAD21^AID^ were differentiated into NPC, and transcription changes were quantified using EU-seq. Transcription changes are compared in cells without or with depletion RAD21^AID^ by treatment with IAA (500uM), and after A-485 treatment (1h). Regulated genes were categorized into indicated class based on the indicated fold-change in gene expression. The data are from 12 biological replicates. **(B)** Number of genes regulated in NPC after 1, 2, and 4 hours of RAD21^AID^ depletion and 1 hour of A-485 treatment (left panel). Fraction of genes regulated in NPC after 1, 2, and 4 hours of RAD21^AID^ depletion and 1 hour of A-485 treatment (right panel) **(C)** Overlap between genes regulated after 1, 2, and 4 hours of RAD21^AID^ depletion in NPC. Number of regulated genes is shown. **(D)** Fraction of RAD21^AID^ regulated genes in NPC that are bound by CTCF and RAD21 within the specified distance from the regulated gene TSS.

**Supplemental Figure 9.**
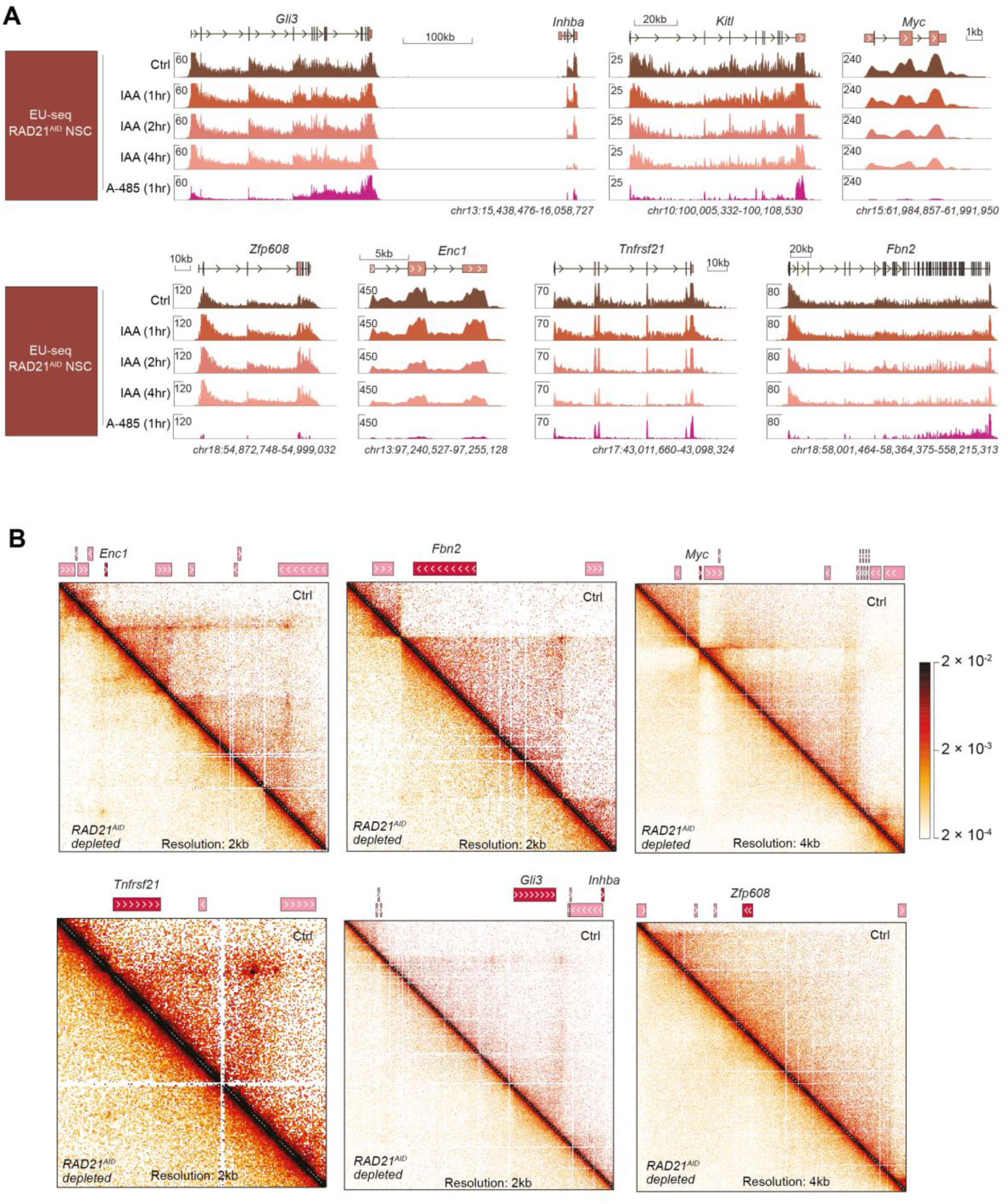
RAD21^AID^ depletion causes gene-selective transcription downregulation in NPC, related to Figure 7. **(A)** Genome browser tracks showing downregulation of known (*Fbn2*, *Myc, Gli3*, *Zfp608*) and putative (*Inhba*, *Tnfrsf21*) enhancer targets by RAD21^AID^ depletion and A-485 treatment in NPC. Some of these targets (*Fbn2*, *Myc, Enc1, Tnfrsf21*) are downregulated both by RAD21^AID^ depletion and A-485 treatment, but others (*Gli3, Zfp608*) are only downregulated by A-485. Of note, the *Gli3* proximal gene, *Inhba*, is downregulated by both RAD21^AID^ depletion and A-485 treatment. **(B)** Micro-C detected chromatin interactions in the indicated loci in wild-type mESC, and change in loop strength after RAD21^AID^ depletion. Micro-C data are from reference ^20^. In several RAD21-regulated genes in NPC (*Enc*1, *Fbn2*, *Myc, Gli3*, *Tnfrsf21*), chromatin loops are visible in mESC but not apparent in others (*Zfp608*).

## Discussion

The roles of cohesin and CTCF in gene regulation are extensively studied. Yet, CTCF’s historical functions as a transcription activator or repressor remain doubted ^63^, the nature of enhancers that rely on cohesin-dependent looping is still not well defined, and the scope of cohesin-CTCF-mediated looping in global gene regulation remains a matter of ongoing discussion ^20–22,43,51,64^. By analyzing the architectural and non-architectural roles of cohesin and CTCF in the same system (**Figure 7D**), this work introduces four major conceptual advances to these ongoing discussions: (1) The scope of cohesin and CTCF in mammalian gene regulation is much broader than previously appreciated. (2) Only a single enhancer type appears to use cohesin-dependent looping for gene activation. (3) Independently of its architectural roles, CTCF functions as a position and orientation-specific transcription repressor and activator, controlling the expression of housekeeping genes, including ones that are pan-essential for mammalian cell proliferation. (4) CTCF and CTCFL share transcription activation ability but differ in their ability to anchor cohesin loops. <u>To the best of our knowledge, these specific conclusions presented have</u> <u>not been published in any prior publications.</u> Below, we discuss how these findings address long-standing questions and reconcile differing perspectives on the roles of cohesin and CTCF in global gene regulation.

### Cohesin and CTCF have broad but subtle roles in gene activation

This work focused on investigating roles of cohesin and CTCF in gene regulation rather than 3D genome organization, as the latter has been extensively studied. Prior research has already examined cohesin-CTCF-dependent looping using acute CTCF^AID^ and RAD21^AID^ depletion in the same cell line ^20,22^, and loss of cohesin-anchored loops in CTCF^YF-AA^ mutant cells has been demonstrated across different cell types^47,48^.

Our study shows that acute depletion of CTCF and RAD21 in mESCs dysregulates hundreds of genes, revealing a much broader role for cohesin and CTCF in gene regulation than previously appreciated ^20–25^. Earlier studies likely underestimated this scope by excluding weakly regulated genes due to stringent statistical thresholds, which explains the differences in findings. While statistical testing is essential, we argue that fold-change-based classification more accurately captures biological regulation in this context. Though the fold-change approach inevitably includes some false positives, our conclusions are robust and supported by orthogonal assays, including RNA-seq, ChIP-seq, ATAC-seq, and plasmid reporter assays (**Supplemental Note 1**). The unequivocal differences in the position- and orientation-specific CTCF enrichment in genes downregulated by CTCF depletion, along with CBP/p300 dependence in genes downregulated by RAD21 depletion, strongly suggest these changes are likely primary effects rather than secondary ones. Despite this collective evidence, if one dismisses the statistically non-significant genes as noise in quantification and/or secondary effects, we are left with the same old conclusion: acute cohesin and CTCF depletion cause regulation of very few genes.

Interestingly, the regulatory scope of cohesin is even more extensive in NPCs where RAD21^AID^ depletion decreases transcription of as much as half of the most strongly (>4-fold) regulated enhancer candidate targets. This implies that the role of chromatin looping in some differentiated cells may be even more prominent than in pluripotent stem cells. Mammals have hundreds of different cell types and enhancers, and their target genes tend to be highly cell-type-specific. Thus, cohesin-CTCF-dependent looping likely affects different genes across various cell types. Extrapolating from two cell types, we propose that cohesin-dependent looping could influence thousands of genes in mammals, underscoring the broad role of 3D genome organization in gene regulation at an organismal level. However, many candidate enhancer targets remain unaffected in CTCF^YF-AA^ cells where the anchoring of cohesin loops should be permanently weakened, implying that cohesin-CTCF loops may not be essential for all E-P interactions.

One explanation is that additional factors contribute to E-P looping ^61,65–67^. Alternatively, cohesin and CTCF might establish initial E-P interactions that are then maintained by a “molecular memory” following their depletion ^20,43,68^. However, the nature of molecular memory and its lifetime remain unknown. Inspired by Paul Nursés call for biologists to generate both new data and ideas ^69^, we propose an alternative model to stimulate discussion on this topic. We posit that architectural proteins mediate some E-P interactions, but this is not a universal requirement for all enhancers. Instead, proteins that bind in enhancers and promoters, such as transcription factors, chromatin remodelers, and coactivators, may mediate shorter-range, transient E-P interactions. This idea is supported by recent findings that transcription factors and chromatin remodelers can independently drive higher-order chromatin organization in yeast without architectural proteins ^70^. However, the likelihood of such E-P interactions would decrease with increased distance. For genes distal enhancers, chromatin looping, for example, by cohesin and CTCF, can decrease effective E-P distances and promote dedicated long-range interactions that would otherwise not occur. This model could explain why not all enhancer-regulated genes are regulated by cohesin and CTCF depletion. This concept could also explain the striking genome-scale association between the distance and strength of CBP/p300-dependent enhancers and their dependency on CBP/p300 activity ^38^. Furthermore, the model could explain why the synthetic insertion of an enhancer near non-enhancer-regulated genes in the native context can effectively boost the transcription of proximal genes ^37^.

### Cohesin facilitates gene activation by CBP/p300-dependent enhancers

The predominant model in the field is that cohesin regulates transcription by chromatin looping ^1–5^. Other models suggest that cohesin regulates transcription by recruiting transcription factors ^71–73^ and counteracting polycomb-mediated gene repression ^23^. We find that: (1) RAD21^down^ shows enrichment of enhancers in their proximity. (2) RAD21^down^ are significantly biased for regulation by CTCF. (3) RAD21^down^ are strongly biased for regulation by CBP/p300, and the cohesin loop anchoring-deficient CTCF^YF-AA^ has selectively lost the ability to activate CTCF-dependent genes that require CBP/p300. (4) RAD21-regulated genes include several bonafide enhancer targets that are shown to require cohesin-mediated looping. These results are consistent with the looping model.

If the main role of cohesin was to recruit transcription factors, it would imply that cohesin selectively facilitates transcription factor recruitment to a subset of regulatory elements that function by using CBP/p300. In mouse thymocytes, genetic deletion of *Rad21* preferentially downregulates enhancer proximal genes, but H3K27ac, H3K4me1, and transcription in enhancers remain unaffected ^74^, suggesting a limited role of cohesin in recruiting transcription factors. Also, the abundance of polycomb-catalyzed H3K27me3 in RAD21^down^ genes appears comparable to genes not regulated by RAD21. Although we cannot completely rule out the non-architectural roles of cohesin, evidence strongly suggests that its primary function in gene regulation involves promoting looping between enhancers and promoters.

Notably, among the genes downregulated by more than twofold after 2-4 hours of RAD21^AID^ depletion, 92-100% are activated by CBP/p300 (**Figures 1D**, **7C**). Genes weakly downregulated by RAD21^AID^ depletion show somewhat lower overlap with CBP/p300 targets, likely due to quantification noise, indirect effects, or false positives. Based on this observation, we suggest that cohesin may almost exclusively promote interaction with CBP/p300-dependent enhancers. Future research should investigate how other enhancer types, acting without CBP/p300, engage with their distal targets. Those enhancers may use other architectural proteins that can promote E-P interactions independently of cohesin and CTCF, such as YY1 and LDB1 ^65–67^.

### Coming full circle: CTCF is more than a looper

Originally identified as a transcriptional activator and repressor ^49,59,75,76^, CTCF binding in the TSS downstream region inhibited transcription from the *c-myc* promoter, whereas binding in the TSS upstream region activated transcription at the *App* promoter ^49,59^. However, these early conclusions were derived from plasmid-based reporters and focused on a few promoters, so the relevance of these findings to in vivo gene regulation remained uncertain. With the discovery of CTCF’s role in 3D genome organization, it was proposed that CTCF’s traditional regulatory functions—transcriptional activation, repression, insulation, and imprinting—may all be secondary to its primary role as a genome-wide organizer of chromatin architecture ^63^. Supporting this model, deletion or inversion of CTCF binding sites or inverting their orientation can rewire E-P loops and alter gene expression ^5–9^, and orientation-specific binding of CTCF in promoters is essential for activation of dozens of protocadherin genes ^7,10,77^.

Our findings confirm many prior observations and provide new insights by disentangling the cohesin-dependent and independent roles of CTCF in gene activation. Among the diverse mechanisms implicated in CTCF-dependent gene regulation ^14,49,51,59,76,78–81^, work suggests five mechanisms through which CTCF regulates the expression of hundreds of genes in vivo. (1) CTCF facilitates the activation of enhancer targets by anchoring cohesin loops. (2) CTCF prevents inadvertent gene activation by blocking enhancer mistargeting. (3) CTCF enforces transcription fidelity at bidirectional promoters by dampening anti-sense transcription. (4) CTCF acts as a canonical transcription activator. (5) CTCF possibly represses transcription in very few instances. A recent study suggests that other housekeeping transcription factors, such as NRF1 and SP1, share this position-dependent activator and repressor function principle of CTCF ^50^.

The delineation of CTCF roles underscore that CTCF’s architectural function is crucial for cell-type-specific gene expression through enhancers, whereas its non-architectural role primarily regulates housekeeping genes. In CTCF^AID^-depleted cells, most strongly downregulated genes are activated by promoter-bound CTCF independently of cohesin or enhancers. This non-architectural function explains why genes downregulated by CTCF^AID^ and RAD21^AID^ depletion overlap only partially ^20,31^ and why genes downregulated by CTCF^AID^ depletion show a stronger bias for CTCF binding in their promoters. These findings reconcile historical and contemporary models of CTCF-dependent gene regulation, showing that CTCF is more than just a “looper ^63^.”

Of note, the bias for forward-oriented CTCF binding in promoters of genes downregulated after CTCF depletion has been known for a decade, but the reason for this bias has not been investigated and the results were mostly interpreted in the context of enhancers and 3D genome organization ^20,22,25,31,43–45^. The enrichment of CTCF in promoter proximal regions is explained by the role of CTCF in facilitating interaction with distal enhancers ^10–12^ ^13^. Consequently, latest reviews discuss CTCF’s roles in gene regulation almost exclusively in the context of 3D genome organization ^43^. <u>Our conclusion is different</u> <u>and adds a new dimention to the current models</u>. We show that, distinct from its architectural role, promoter-bound CTCF functions as transcription activator. We are unware of any prior report demontrating a non-architectural, looping-independent function of promoter-bound endogenous CTCF in activating housekeeping genes in native chromatin.

### CTCF is a position-specific activator and repressor of sense and anti-sense transcription

The functional diversity of CTCF appears to be dictated by the position and polarity of its DNA binding (**Figure 7D**). CTCF acts as a transcription activator only when bound in a forward orientation within a specific range upstream of a core promoter. If CTCF binds outside this narrow window, either further upstream or downstream, it assumes roles as a repressor or non-activator. In distal regions, CTCF typically functions as a chromatin organizer, promoting E-P interactions or blocking enhancers from mistargeting genes. This functional diversity and the positional and orientational dependency of CTCF appear unmatched by any other transcription regulator.

This raises intriguing questions about how these binding positions lead to such distinct outcomes. While the intricacies remain to be resolved, position-specific CTCF binding appears to distinctly affect promoter accessibility, Pol II binding, and cohesin loop anchoring. Although CTCF generally does not determine promoter accessibility, it is essential for accessibility and proper Pol II recruitment at certain promoters where it functions as an activator. Conversely, when CTCF acts as a repressor, it seems to hinder Pol II binding or positioning—potentially by obstructing factor binding required for pre-initiation complex assembly or impeding Pol II elongation.

Further, the positioning of CTCF near the promoter appears to differentiate its impact on sense and antisense transcription. When CTCF binds upstream of the promoter, it prevents antisense transcription, while binding within or just downstream of the core promoter represses sense transcription. This implies that CTCF represses sense and antisense transcription by the same principle—affecting Pol II binding or elongation—with its impact depending on its binding position relative to the TSS. In this respect, CTCF appears to act as a “wheel choke” that influences Pol II binding or elongation, depending on its position.

The positional and orientation bias of CTCF at promoters of CTCF-dependent genes suggests that CTCF directly binds in promoters, which is distinct from the previously proposed model of indirect CTCF recruitment through interactions with TFII-I ^82^. These findings reveal how CTCF’s regulatory versatility stems from its spatial and directional binding preferences, making it both a unique architectural organizer and a context-dependent transcriptional regulator.

### Functional divergence of *Drosophila* and vertebrate CTCF

CTCF is evolutionarily conserved between *Drosophila* and vertebrates, showing functional similarities and notable differences. For example, CTCF functions as an enhancer blocker both in *Drosophila* and mammals ^14–16,35^. However, unlike mammalian CTCF, *Drosophila* CTCF neither functions as a cohesin loop anchor nor is required for embryonic development or the proliferation of cultured *Drosophila* cells ^32,34,83,84^. Additionally, unlike in mammalian cells, genes downregulated in CTCF knockout *Drosophila* exhibit only a modest bias for CTCF binding in their promoters ^32,34^. These differences suggest that while CTCF’s enhancer-blocking function is ancient, its roles as a cohesin loop anchor and transcription activator likely evolved later in vertebrates.

The importance of CTCF’s vertebrate-specific, cohesin-loop anchoring function is highlighted by extensive gene dysregulation in CTCF^YF-AA^ expressing cells, while the importance of transcription activator function is highlighted by downregulation of hundreds of genes after the depletion of CTCF^YF-AA^. Notably, CTCF’s transcription activator function controls the expression of more than a dozen common essential genes, suggesting that CTCF’s pan-essentiality for mammalian cell proliferation likely stems from its role as a transcription activator. This may explain the distinct requirement for CTCF in cell proliferation in *Drosophila* versus mammals. Given that CTCF activates housekeeping genes, which are often highly conserved, it is tempting to postulate whether its transcription activator role arose before its cohesin-anchoring role.

This work also clarifies the divergent functions of CTCF and its paralog CTCFL. CTCFL overexpression can activate the transcription of CTCF^down^ genes bound by CTCF in their promoters, but CTCFL cannot anchor cohesin loops. This implies that aberrant expression of CTCFL in cancers may interfere with CTCF-dependent genome architecture ^85^, while preserving the activation of cell proliferation essential genes directly activated by promoter-bound CTCF. This may explain why CTCFL, unlike CTCF, preferentially binds promoters over distal regulatory regions ^55,56^.

Collectively, by revealing the roles of cohesin and CTCF in regulation in vivo gene regulation and disentangling their architectural and non-architectural roles, our findings provide an integrated model of gene expression control by cohesin, CTCF and enhancers, with important implications for understanding gene regulation in physiology and disease.

**Supplemental Note 1**

Most genes regulated after CTCF^AID^ and RAD21^AID^ depletion do not pass statistical thresholds. The following evidence supports that weak transcription changes observed in our analyses are non-random and reflect genuine perturbation-induced changes:

1. CTCF-regulated genes identified in EU-seq data are consistently regulated in independently published RNA-seq data in the same cell line (**Supplemental Figure 1D**).
2. Most RAD21^down^ identified in initial experiments show consistent downregulation in additional replicates (**Supplemental Figure 2C**).
3. Genes down-regulated by CTCF and RAD21 depletion strongly differ in their regulation by A-485 (**Figure 1**).
4. Among CTCF^down^ genes, those not downregulated by A-485 are strongly bound by CTCF at their promoters, while those downregulated by A-485 show no strong CTCF binding (**Figure 3C-D**).
5. Genes regulated by CTCF and RAD21 show notable differences in cell-type specificity (**Figure 4A**). Within CTCF-downregulated genes, those requiring CTCF-anchored loops and those not requiring loops show opposite cell-type-specific expression patterns.
6. Acute depletion of CTCF^YF-AA^ selectively decreases the transcription of CTCF-dependent genes bound by CTCF at their promoters (**Figure 4B-D**).
7. CTCF^down^ genes exhibit strong polarity in CTCF binding at their promoters (**Figure 4E**).
8. CTCF^down^A-485^ND^ genes exhibit strong positional binding of CTCF within a narrow, TSS upstream window (**Figure 4G**).
9. The loop anchoring-defective CTCF^YF-AA^ mutant activates promoters of several CTCF^down^ genes in reporter assays (**Figure 5A**).
10. Overexpression of BORIS selectively restores transcription of a subset of CTCF^down^ genes bound by CTCF at their promoters (**Figure 5B**).
11. CTCF-dependent transcription changes in EU-seq strongly correlate with CTCF-dependent promoter chromatin accessibility in ATAC-seq, analyzed using an independently generated CTCF depletion cell line (**Figure 6A-C**).
12. CTCF-dependent transcription changes in EU-seq strongly correlate with CTCF-dependent recruitment of Pol II in ChIP-seq, again measured in an independently generated cell line (**Figure 6D-E**).
13. Like mESC (**Figure 1**), RAD21^AID^ depletion in NPC causes preferential downregulation of CBP/p300-dependent genes (**Figure 7**).

These results in independently generated cell lines and different assays (EU-seq, RNA-seq, ATAC-seq, ChIP-seq, and plasmid reporters) support the presented conclusions.

**Supplemental Note 2**

It is suggested that the removal of cohesin results in gene downregulation through increased polycomb-mediated repression ^23^. To investigate whether gene downregulation after RAD21^AID^ depletion results from reduced enhancer targeting or increased polycomb interaction, we examined polycomb-catalyzed H3K27me3. H3K27me3 is more frequently enriched in low (TPM 5-15) expressed as compared to high (TPM >15) expressed genes, and genes regulated by A-485, CTCF, and RAD21 show slightly more H3K27me3 enrichment than not regulated genes (**Supplemental Figure 2c**). Nevertheless, at the used expression threshold (TPM >15), only ∼15% of RAD21^down^ genes are marked by H3K27me3, which is comparable to H3K27me3-positive A-485^down^ genes.

## Acknowledgments

We thank the members of the Choudhary lab for their helpful discussions. The Novo Nordisk Foundation Center for Protein Research is financially supported by the Novo Nordisk Foundation (NNF14CC0001). C.C. is supported by the Novo Nordisk Foundation Distinguished Investigator Bioscience and Basic Biomedicine (NNF22OC0074677). We thank the CPR Imaging Platform, the CPR Big Data Management Platform and the CPR and DanStem Genomics Platform for their assistance. We also thank Dr. Rob Klose for providing mESC expressing *Rad21-mAid-Gfp* (RAD21^AID^), and Drs. Nora Elphege and Benoit G Bruneau for providing mESC expressing *Ctcf-mAid-Gfp* (CTCF^AID^).

## Materials and methods

### Cell culture

We used previously published mESCs expressing degron-fused *Rad21-mAid-Gfp* (RAD21^AID^) ^23^ and *Ctcf-mAid-Gfp* (CTCF^AID^) ^22^. RAD21^AID^ expressing NPCs were generated by differentiating RAD21^AID^ mESC ^23^. *GFP-FKBP12^V36F^-Ctcf* and *GFP-FKBP12^V36F^-Ctcf^Y22A^*^6^*^/F^*^228^*^A^* cells were generated using mESC (ES-E14TG2a) from European Collection of Authenticated Cell Cultures (ECACC) (Sigma-Aldrich; 08021401).

Unless indicated otherwise, mESCs were cultured in a custom-made (C.C.Pro GmbH) N2B27 medium consisting of a 1:1 mix of DMEM/F12 and neurobasal medium but lacking arginine and lysine. Before use, the medium was supplemented with 1,000 U ml−1 leukemia inhibitory factor (Merck Millipore), 1 µM PD0325901, 3 µM CT-99021 (custom-made by ABCR GmbH), 100 µM 2-mercaptoethanol, 0.5× B27 supplement (Thermo Fisher Scientific), 0.5× N2 supplement (made in house or from Thermo Fisher Scientific), L-lysine and L-arginine. Unless indicated otherwise, the cells were treated with the following chemical concentrations: A-485 10μM, dTAG-13 0.1 μM, or IAA 500 μM.

### Differentiation of RAD21^AID^ mESC to NPC

RAD21^AID^ expressing mESC were first differentiated into EpiSCs by culturing them on dishes coated with 10ng/ml fibronectin (Merck Millipore), in N2B27 supplemented with 20ng/ml Activin A and 12ng/ml Fibroblast growth factor 2 (FGF2) (PeproTech). EpiSCs were further differentiated into NPC by seeding 0.5-1×105 cells per cm2 to fibronectin-coated dishes in N2B27 without growth factors. After 5 days, cells were dissociated with 0.5x TrypLE (Thermo Fisher Scientific) and seeded to 0.1% gelatinized tissue culture plates in N2B27 supplemented with 10ng/ml FGF2 and 10ng/ml epithelial growth factor (EGF) (PeproTech). NPCs were cultured in 0.1% gelatin-coated plates in N2B27 supplemented with 10ng/ml FGF2 and 10ng/ml EGF.

### CRISPR-Cas9 gene editing

*Puro-P2A-Gfp-Fkbp^12F36V^*-*Ctcf* homology arm or *Puro-P2A-Gfp-Fkbp^12F36V^*-*Ctcf^Y22A^*^6^*^/F^*^228^*^A^* homology arm were introduced into the endogenous CTCF locus using the CRISPR–Cas9 system (full sequences of the targeting constructs are provided in the **Supplementary Table1**). mESC were grown in 2i media and 500ng of Donor and 1μg of sgRNA plasmid were transfected using Lipofectamin 2000 (Thermo Scientific Fischer, 11668019), according to the manufacturer’s instruction. After overnight incubation, the transfection mix was exchanged to normal 2i media. The next day, all cells were dissociated using TrypLE (Thermo Fisher Scientific), spun down, transferred to a new plate, and selected using 0.5 μg/ml Puromycin (Invitrogen, ant-pr-1). After six days of selection, the cells were transferred to normal 2i media, and individual colonies were picked and genotyped using KOD Xtreme Hot Start DNA Polymerase (Novagen, 71975).

### ATAC-seq

CTCF-dependent chromatin accessibility was analyzed in mESC in which endogenous CTCF was fused with a green fluorescent protein and the degron tag FKBL12^F36V^ (*GFP-FKBP12^V36F^-Ctcf*) cells. Cells were first washed in pre-warmed PBS and dissociated from the plate using TrypLE (Thermo Fisher Scientific). After disaggregating the cells, cold media was added to stop the trypsinization. 2X10^5^ cells were transferred to LoBind microfuge tube (Eppendorf) and centrifuged at 500 g for 5min at 4°C.

After centrifugation, the cells were resuspended in buffer A (10mM HEPES pH 7.9, 10mM KCl, 1.5mM MgCl2, 0.34M Sucrose, 10% Glycerol, 10% Triton-X), mixed by tapping and incubated on ice for 7min. Following another centrifugation, nuclei were resuspended in lysis buffer (10mM Tris-HCl pH7.4, 10mM NaCl, 3mM MgCl2, 0.3% Igepal). A quarter of nuclei were then transferred to a new Lobind tube, briefly vortexed, incubated on ice for 15 min, vortexed again briefly, and pelleted down at 600 g for 10 min at 4°C. After removing the lysis buffer, nuclei were recovered with 2.5 μL of 2X TD buffer and 2.5 μL transposase (TDE1) mixture (Illumina). Following mixing by pipetting, the nuclei were incubated for 30 min at 37°C on a rocking mixer. The transposed DNA was purified using the MinElute PCR Purification kit (QIAGEN) following the manufacturer’s protocol. The transposed genomic regions were PCR-amplified using Nextera Primers (https://www.encodeproject.org/documents/860c95a8-fdc4-46bd-b9b5-d44242ff20ef/@@download/attachment/ATAC_garberlab_protocol.pdf) and NEBNext High-Fidelity 2x PCR Master Mix (NEB, M0541). The PCR product was purified using Agencourt® AMPure® XP beads (Beckman Coulter) and sequenced on a NextSeq 500 sequencer (Illumina).

### ChIP-seq

The cells were cross-linked for 10 min at room temperature with 1% formaldehyde solution with gentle agitation and quenched with 0.125 M glycine. Cross-linked cells were washed once with PBS, added swelling buffer (10 mM Tris-HCl pH8.0, 1.5 mM MgCl2, 10 mM NaCl, 0.5% NP-40), and incubated on ice for 10 min. After centrifugation, pellets were suspended in RIPA buffer (10 mM Tris-HCl pH8.0, 1 mM EDTA, 140 mM NaCl, 1% Triton X-100, 0.1% SDS, 0.1% Sodium Deoxycholate) containing protease inhibitor cocktail (Roche), and then were sonicated by using a BioRuptor sonicator (Diagenode). After centrifugation, solubilized chromatin was recovered and incubated with 5 μg of Pol II antibody (#14958, Cell Signaling Technology) bound to Dynabeads M-280 sheep anti-Rabbit IgG (Thermo Fisher Scientific). After overnight incubation at 4°C, the magnetic beads were washed with RIPA buffer twice, RIPA high salt buffer (10 mM Tris-HCl pH8.0, 1 mM EDTA, 500 mM NaCl, 1% Triton X-100, 0.1% SDS, 0.1% Sodium Deoxycholate), LiCl buffer (10 mM Tris-HCl pH8.0, 1 mM EDTA, 250 mM LiCl, 0.5% NP-40, 0.5% Sodium Deoxycholate) and TE buffer (Invitrogen). The bound materials were eluted with elution buffer (10 mM Tris-HCl pH8.0, 5 mM EDTA, 300 mM NaCl, 0.1% SDS) overnight at 65°C and treated with RNaseA for 30 min for 37°C, followed by further incubation with Proteinase K for 1 hour at 37°C. Then DNA was purified with a PCR purification kit (QIAGEN). ChIP-seq libraries were prepared by using the NEBNext Ultra II DNA prep kit (NEB E7645) following the manufacturer’s instruction and sequenced on a NextSeq 500 sequencer (Illumina).

### EU-seq

To analyze nascent transcription changes under various conditions, cells were treated with DMSO or with the indicated chemicals for the specified time points. Unless indicated otherwise, A-485 at 10μM, dTAG13 0.1 μM, or IAA at 500 μM were used for treatment. Cells treated with or without chemicals were pulse-labeled with 0.5 mM 5-EU for 20 min before sample collection. After the 5-EU labeling, cells were quickly washed once with PBS, and immediately lysed in RLT buffer, and RNA was isolated using the RNeasy Mini Kit (QIAGEN) according to the manufacturer’s protocol. Labeled RNA was further enriched using the Click-iT® Nascent RNA Capture Kit (Thermo Fisher Scientific) according to the manufacturer’s instructions. EU-seq libraries were prepared using the NEBNext® Ultra II RNA First Strand Synthesis Module (E7771) combined with NEBNext® Ultra II Non-Directional RNA Second Strand Synthesis Module (E6111) or NEBNext® Ultra II Directional RNA Library prep kit for Illumina (E6111), following the manufacturer’s instructions and subsequently sequenced on either NextSeq 500 sequencer or NextSeq 2000 sequencer (Illumina).

### Cloning of CTCF promoter luciferase plasmids

5µg pGL421 MLL promoter plasmid (Addgene: 118695) was linearized by digestion with XhoI and HindIII and gel extraction purified. Genomic sequences encompassing promoters of analyzed genes were amplified by nested PCR using first KOD polymerase and then PrimeStar MAX DNA polymerase to generate homology regions to the linearized pGL421. Pieces were gel extraction purified and inserted into the vector by Hot Fusion cloning followed by transformation, clone picking, plasmid isolation, and sequence verification by Sanger sequencing. Where indicated, promoter luciferase plasmids with inverted and mutated CTCF sites were generated by using the plasmids with the native promoters as template for amplification of two pieces upstream and downstream of the putative CTCF site with primers in between encoding the inversion/mutation as well as homology for homology-directed cloning. The two pieces were combined with linearized pGL421 and cloned by Hot Fusion cloning, followed by clone picking, plasmid isolation, and sequence verification by Sanger sequencing. Promoter sequences used for reporter assays are provided in **Supplementary Table 2**.

### CTCF promoter luciferase assay

Promoter reporter activity was measured using the Dual-Glo® Luciferase Assay System (https://www.semanticscholar.org/paper/Dual-Luciferase-TM-Reporter-Assay%3A-An-Advanced-and-Sherf-Navarro/a798b2a51acc2a146bbeb4705a4168f15baf3b6f). Briefly, 500ng pGL474 and 1000ng pGL421 in 150µL OptiMEM with 3µL Lipofectamine 2000 were incubated for complex formation at RT 10 min and transfected to 5e5 CTCF^YF-AA^ mESC for 10 min. Cells were collected by centrifugation, and the transfection mix was removed, followed by resuspension in 1.5mL 2i LIF medium. 150µL (5e4) of the cells were seeded to 4 wells for each transfection in a geltrex-coated 96-well plate and allowed to settle 20min at room temperature before incubating overnight in the incubator for attachment to the surface. The next day, 18h after transfection, treatment of the cells with DMSO (1:500) or dTAG-13 (200nM) was started and allowed to proceed for 6h. Then, cells were washed once with PBS at RT and incubated with 48µL PBS. Cells were lysed, and firefly luciferase substrate was added with 48µL of Dual Glo Luciferase assay reagent to the plate, followed by 10 min incubation at RT and bioluminescence measured with a Tecan Spark using default settings. Then 48µL Dual Glo Stop & Glo reagent was added to the plate to quench firefly luciferase activity and provide the substrate for Renilla luciferase. Following 10 min incubation at room temperature, bioluminescence was measured. The luminescence values for Firefly and Renilla luciferase were adjusted by subtracting the average luminescence values of the blank wells on the same plate. To mitigate the high variability associated with low luminescence levels, any subtracted luminescence value less than 30 relative light units (RLU) was adjusted to 30. Relative response ratios (rrr) were then calculated as the ratio of the blank-corrected luminescence value of Firefly luciferase to that of Renilla luciferase. These ratios were further normalized by the rrr value of the wild-type promoter treated with DMSO (normalized rrr).

### Immunoblotting

The day prior to sample preparation, 5e5 cells were seeded for each sample in a 6-well plate. The next day, treatment was done with vehicle control (DMSO, 1:500) or dTAG-13 (100nM) for 2h. Cells were collected by scraping and lysed in RIPA followed by 5×10s sonication at 40% amplitude and centrifugation 10min at 21000g to collect the sample from the supernatant. The protein concentration was determined with the bicinchoninic acid (BCA) assay. 25µg protein sample mixed with 4xNuPage LDS to 1x was heated at 70°C for 5 min, and the sample was loaded into a 4-12% NuPage Bis-Tris 1.0mm mini protein gel.

Seeblue Plus2 pre-stained was used as a ladder. The gel was run with 1xMOPS buffer 20min at 80V and then 40 min at 200V. Proteins on the gel were transferred to a nitrocellulose membrane in a transfer buffer for 90min at 30V. The membrane was washed with water, stained with Ponceau for 5 min, and washed with water 5min followed by imaging. Ponceau was washed away with 0.1M NaOH for 2 min, and the membrane was washed with water, followed by overnight blocking with 5% skim milk in 1X TBST at 4°C for 14-18h. Milk was washed away with 1xTBST, and the blot incubated with 1:1000 anti-CTCF antibody (rabbit, D31H2, Cell Signaling Technologies) in 1xTBST with 1% BSA for 2h followed by 4×3min washes with 1xTBST. The blot was then incubated with anti-rabbit HRP antibody for 1h, washed 4×3min with 1xTBST, developed with Clarity Western ECL substrate, and imaged on an ImageQuant LAS 4000 or an Amersham ImageQuant 800 system.

### Immunofluorescence staining

Treatment, fixation and staining for CTCF microscopy and QIBC were performed as follows. Cells (25000/well) were seeded to geltrex-coated wells in a Perkin Elmer Cell Carrier 96w plate and allowed to adhere overnight. Cells were treated with vehicle control (DMSO, 1:500) or dTAG-13 (100nM) for 2h. Then cell media was removed by inversion of the plate, and the cells were fixed with 4% formaldehyde in PBS for 15min. Cells were then washed with PBS, and the plate was stored in the fridge until staining. For staining, cells were permeabilized with 0.5% triton X-100 in PBS for 5min and blocked 30min with antibody diluent (DMEM with 10% FCS, 0.02% sodium azide, filtered). Then, cells were incubated with anti-CTCF (1:1000) in an antibody diluent for 2h, washed twice with PBS, and incubated with anti-rabbit Alexa 568 in an antibody diluent with DAPI. Finally, the cells were washed twice with PBS. Images of the stained cells were acquired with the screening microscope described below.

### Microscopy and image-based cytometry

Microscopy images and images for image-based cytometry were acquired on an automated Perkin Elmer Phenix spinning disk screening microscope. The microscope was equipped with two 16-bit OEM Andor Zyla sCMOS cameras with 2160×2160 pixels with a 6.5µm pixel size. The objectives used were a 20x water immersion NA 1.0 W Plan apochromat or a 40x water immersion NA 1.1 LD C-Apochromat objective. Harmony 4.8 image analysis software was used for the quantification of images. The analysis pipeline included background correction, nuclear segmentation, filtering of border objects, quantification of the individual signals (mean, sum, and median per nuclei), and generation of the data tables. The data tables were further processed in R using ggplot2 for visualization. Representative example image projections from each condition with set contrast and brightness settings for each staining are displayed.

### Data analysis

#### Reference genome annotation

Mouse and human genome annotation and reference genome were downloaded from the GENCODE (mouse; GRCm38 release 25, human; Grch37 release 29)^86^. As a reference, the transcript type of “protein coding” and “lincRNA” was chosen as representative genes. To build the mESC transcripts reference used in the EU-seq, we applied two-step transcripts annotation. First, the expressed transcripts were selected using the criteria of transcripts per million transcripts (TPM) > 2 in RNA-seq and the presence of H3K4me3 ChIP seq peak within 1Kb from TSS. The longest isoform was selected if a single H3K4me3 peak is proximal to multiple transcripts annotated to the same. For the genes where the H3K4me3 peak was not detected, the longest isoform was chosen as a representative transcript and combined with the transcripts reference.

#### Processing RNA-seq data

Adaptors were trimmed using Cutadapt (https://doi.org/10.14806/ej.17.1.200). Reads were mapped to the reference genome using STAR (version 2.6.1a)^87^ with removing the non-canonical junctions. Reads mapped to tRNA and rRNA regions were removed using Bedtools (version 2.23)^88^. The number of reads mapped to an exon was counted on a gene basis by using HTseq (version 0.11.1)^89^. The TPM was calculated using R.

#### Processing EU-seq data

Adaptor trimming was performed as described in the RNA-seq section. Read sequences were aligned to the mm10 mouse genome using bwa meme (version 1.0.4) ^90^ with the soft clipping option for supplementary alignments. Low-mapping quality reads ( < q 10) were removed using samtools (version 1.4)^91^. Reads mapped to rRNA and tRNA regions, obtained from the UCSC genome browser, were removed using Bedtools. For the 30, 60, and 120-min treatment data, based on the Pol II elongation rates, gene body regions of up to 30, 90, and 240 Kb from the TSS were used for differential gene expression (DGE) analysis. The number of reads mapped to defined regions was counted using HTseq. Transcripts with low mapped reads (average reads ≤ 20 between replicates in both control and treatment conditions) were excluded from DGE analysis. Log2 fold-change (FC) was calculated with the default scaling method of DESeq2^92^, using the median of relative abundance. Asymmetrical DGE profiles resulted in a bias towards the opposite direction of the asymmetry. To adjust for this, the median log2 FC value was subtracted from the DEseq output. Unless described otherwise, fold changes from distinct timepoints were summarized by calculating the averages. The TPM was calculated using the filtered transcripts based on read counts. Further, to keep only transcripts with robust expression, those with average TPM of control and treatment conditions >15 were defined as expressed. In the calculation of antisense transcripts regulation, only the strand-specific EU-seq data was used. Reads mapped to the antisense strand of upstream regions of -1.7 kb - -200b from TSS were counted. The regulation was calculated only in antisense transcripts corresponding to expressed sense transcripts, and those with low mapping reads (average reads ≤ 10 between replicates in both conditions) were excluded from DGE analysis. Normalization and asymmetrical bias correction were performed using the scaling factors and median log2 FC values derived from the sense transcripts DGE analysis. For the gene track visualization, we used deeptools and bigWigMerge (UCSC tool)^93^ to generate bigwig files using scaling factors calculated from DESeq2.

#### Classification of EU-seq regulation class

Unless specified otherwise, transcripts in each experimental condition were classified according to their fold changes after treatment, as follows. HU: ≥2-fold up-regulation. IU: <2- and ≥1.5-fold up-regulation. SU: <1.5- and ≥1.3-fold up-regulation. NC: <1.2- fold up-regulation and <1.2-fold down-regulation. SD: ≤1.5- and >1.3-fold down-regulation. ID: ≤2- and >1.5-fold down-regulation. HD: > 2-fold down-regulation. Others: <1.3- and ≥1.2-fold up-regulation, or ≤1.3- and >1.2-fold down-regulation. For the A-485 regulation class, the following criteria were applied. ND: ≤ 1.2-fold down-regulation. ID: ≤2- and >1.5-fold down-regulation. HD1: ≤ 4- and >2-fold down-regulation. HD2: >4-fold down-regulation. For RAD21-regulated genes in NPC, the fold change values at 2hour and 4 hour timpoints are averaged and classified as follows; HD: >1.3 fold downr-regulation on both 2hour- and 4hour-treatment with an average fold changes of >2-fold down-regulation, ID: >1.3 fold downr-regulation on both 2hour- and 4hour-treatment with an average fold changes of ≤2- and >1.5-fold down-regulation, SD: >1.3 fold downr-regulation at 2hour- and 4hour-treatment with an average fold changes of ≤1.5- and >1.3-fold down-regulation. If the number of genes within any class is fewer than 10, the class is combined with the adjacent class, and the combined group is shown with a joint name (e.g., HU+IU). In classifying antisense transcripts, only those exhibiting ≥2-fold up-regulation were selected and manually reviewed to confirm genuine upregulation, ensuring the changes were not due to the changes of overlapping or neighboring transcripts. Antisense transcripts that were confirmed as genuinely upregulated through this manual verification were used for presented analyses.

#### Processing of ChIP-seq data

Adaptor trimming was performed as described in the RNA-seq section. Read sequences were aligned to the mm10 mouse or hg19 human genome using bwa meme (version 1.0.4) ^90^ with the soft clipping option for supplementary alignments. Duplicated reads were annotated and removed using Picard toolkit tools (version 2.9.1, “Picard Toolkit.” 2019. Broad Institute, GitHub Repository. https://broadinstitute.github.io/picard/; Broad Institute). For the paired-end reads, only the properly aligned read pairs were retained, using samtools. Peak calling of CTCF, RAD21, H2BK20ac, and H3K27ac, TBP and H3K4me3 ChIP-seq were performed using LanceOtron with the default model (wide-and-deep_jan-2021)^94^. Peak calling of H3K27me3 was performed using epic2 (version 0.0.47) ^95^. In the H2BK20ac and H3K27ac datasets, the peaks proximal within 2kb are merged using Bedtools^96^. Poorly enriched peaks of maximum peak height < 8 reads mapped per million (rpm) were omitted. Peak heights were calculated using bamCompare^97^ with the following parameters (centerReads, minMappingQuality 10, bin size 20bp, smoothing length 400bp, extension of reads to 200bp, rpm normalization, and input rpm value subtracted). The peak summit was defined as the center of the 20b bin at maximum height in each peak region. To identify the precise peak summit position of CTCF, CTCF ChIP-exo peaks were determined using GEM (version 3.4) ^98^ with the following parameters (minimus k-mers 6, maximum k-mers 13, smoothing 3bp, maximum read count on a base position 20). The CTCF ChIP-seq peaks with summit positions within 200b from the center of ChIP -exo peak region were retained and used for further analysis. CTCF peaks were categorized into quartiles Q4, Q3, Q2, Q1 based on the ChIP-seq peak height, from highest to lowest. The CTCF peak summit positions were defined by the center of ChIP-Exo peak regions. EPDnew ^99^ was used to determine the precise TSS position as follows; If the EPD TSS position was located within 200b from the GENCODE TSS position of the corresponding gene, the EPD TSS position was used, otherwise GENCODE TSS position was used. The distance between ChIP-seq peak and TSS indicates the distance between the peak summit and the TSS. For the calculation of Pol II ChIP-seq changes after CTCF degradation, we assumed that Pol II binding remains unaffected at gene body regions of the genes that are not changed (NC) in EU-seq analysis. The scaling factors were calculated by DESeq2 default normalization methods using the Pol II ChIP read counts at gene body regions of NC genes in the CTCF degradation experiments. The aggregate ChIP-seq profiles were plotted using deepTools2 (version 3.5.2) ^97^. For gene track visualization, IGV^100^ (version2.16) was used. For the visualization of ChIP-seq and ATAC-seq profiles on gene track, we used 1bp binned, input-subtracted values without any smoothing (CTCF, TBP, Pol II, and ATAC-seq), or 20bp binned, input-subtracted values with smoothing using proximal 400bp (H3K4me3).

#### Defining enhancer strength

Enhancer regions were defined as overlapping regions of H2BK20ac and H3K27ac peaks. To assess the enhancer strength, H2BK20ac reads were counted within these regions. Reads mapped near TSS (+/-500bp regions) were excluded to avoid the potential contribution of promoter acetylation. The read count was normalized by using reads per million (rpm), and input rpm values were subtracted at the corresponding regions. Peaks with H2BK20ac > 1 rpm were retained and used to define enhancer peaks. These enhancers were then ranked by their H2BK20ac rpm values, and the top 10, 25, 50, and 75% enhancers were classified based on their H2BK20ac enrichment within the identified enhancers.

#### Processing of ATAC-seq data

Adaptor trimming was performed as described in the RNA-seq section. Paired-end read sequences were aligned to mm10 genome using BWA meme (version 1.0.4)^90^ with the soft clipping option for supplementary alignments. Duplicated read pairs were removed using Picard-tools (version 2.9.1, “Picard Toolkit.” 2019. Broad Institute, GitHub Repository. https://broadinstitute.github.io/picard/; Broad Institute). We filtered out low-quality reads with a MAPQ score of less than 10 and non-primary alignments and retained only the properly aligned read pairs using samtools. Peak regions were called from the merged alignment files of both control and dTAG-13-treated samples using LanceOtron and its default model (wide-and-deep_jan-2021)^101^. Peak height was calculated using bamCoverage with the following parameters (centerReads, bin size 20bp, smooth length 400bp, extend reads 200bp, rpm normalization). Poorly-enriched peaks of peak summit height < 8 rpm were filtered out. For the calculation of ATAC accessibility changes, ATAC reads that were mapped in the summit +/- 100bp regions were counted. After filtering out the low-mapped regions (average reads ≤ 20 between replicates in both control and treatment conditions), log2 fold-change (FC) was calculated with the default scaling method of DESeq2^92^, using the median of relative abundance.

#### Analysis of CTCF ChIP-seq data from different human and mouse cell lines and tissues

The publicly available human and mouse CTCF ChIP-seq data were downloaded from the GEO repository and peaks were called as described in the processing ChIP-seq data section. The criteria for selecting datasets were the following: 1. Corresponding input data is registered in the same series, 2. Peak number > 20,000. One dataset was chosen for each tissue /cell line. As a result, 49 mouse and 65 human tissues/cell line CTCF peak data were used for the analysis. In each dataset, the distance between the TSS of “protein_coding” or “lincRNA” transcripts and the nearest peak summit was calculated. If CTCF peak summit position is within 200bp upstream or 50bp downstream of TSS, the corresponding gene was defined as CTCF peak (+).

Each CTCF peak was ranked by the peak height from highest to lowest, and the top 20,000, peaks in each dataset were used for comparison. P-values were calculated using Kolmogorov-Smirnov test and test and Benjamini & Hochberg method was used for multiple comparison correction.

#### CTCF motif analysis

Enriched motifs were detected by comparing the sequences of +/- 50b region from CTCF peak summits with their dinucleotide shuffled sequences. STREME (version 5.5.4) ^102^ was used for motif elicitation by the following parameters (minimum motif length 6, maximum motif length 30, differential motif detection mode) and compared with CTCF (MA0139.1) from JASPAR database (https://jaspar.genereg.net). To summarize the motif occurrences and their orientations, the output motifs from STREME were scanned using the +/- 100b region from CTCF peak summits using fimo (version 5.5.4) ^103^.

#### Gene expression profiles of various tissues in mouse

Cell type-specificity of genes was classified using the FANTOM5 CAGE dataset ^104,105^. In the mouse CAGE dataset, we excluded data from “whole,” “lactating,” and “pregnant” stages. The resultant 151 mouse tissue expression profiles (44 tissues from 11 developing stages) were converted into a binary expression matrix using a gene expression threshold of TPM ≥ 2.

#### Depmap dataset

The processed CRISPR gene effect datasets and common essential gene list were downloaded from Depmap portal (version 23Q2) ^106^. Human orthologues of mouse genes were determined by using the Ensembl Orthologue dataset ^107^ with the following criteria. 1. Homology type is “ortholog_one2one”, or 2. Homology type is either “ortholog_one2many” or “ortholog_many2one” and the confidence score is 1.

#### Statistical analysis

Otherwise mentioned, p-values are calculated using the two-sided Mann–Whitney U test and corrected for multiple comparisons using the Benjamini & Hochberg method (R package stats version 3.6.2).

#### Use of publicly available data

The following publicly available data were used: Mouse and Human gene annotations were from GENCODE (https://www.gencodegenes.org); FANTOM5 CAGE dataset was from ArrayExpress (https://www.ebi.ac.uk/arrayexpress/); Promoter Elements were referenced from EPD (https://epd.epfl.ch//index.php); Motif references were downloaded from JASPAR (https://jaspar.genereg.net/). Depmap datasets were downloaded from the DepMap portal (https://depmap.org). The orthologue reference was downloaded from (https://www.ensembl.org/info/data/biomart).

The following publicly available sequencing datasets were reanalyzed: RNA-seq in ESC (GSE140363). RNA-seq in ESC with or without expressing CTF-CTCFL fusions (GSE140363). EU-seq in ESC treated without and with A-485 (GSE146328). ChIP-exo-seq in ESC on CTCF (GSE98671).

ChIP-seq in ESC on H3K4me3 (GSE146328), TBP (GSE146328), TAF1 (GSE146328), H3K27me3 (GSE186349), Input control (GSE146328), CTCF (GSE178982), Rad21 (GSE178982). Following CTCF ChIP-seq and ChIP-exo-seq data in a variety of tissues and cell lines were reanalyzed. Mouse: 3134 cells (GSE61236), 3T3-L1 cells (GSE95533), AML12 cells(GSE95116), splenic B cells (GSE44637), bone marrow derived macrophages (GSE189975), bone marrow (GSE49847), brain (GSE114606), brown preadipocytes (GSE74189), cerebellum (GSE49847), CFUMk cells(GSE156074), CH12 cells (GSE49847), CMP cells (GSE159503), cortex (GSE49847), distal forelimbs (GSE101714), DP cells (GSE141223), ECOMG-derived neutrophils (GSE93127), embryoid bodies (GSE119874), EpiLC cells (GSE183828), erythroid cells (GSE142006), erythroid progenitor cells (GSE150415), female germline stem cells (GSE137771), forebrain (GSE127870), G1E cells (GSE156074), GMP cells (GSE156074), GSC cells (GSE183828), GSCLC cells (GSE183828), heart (GSE91813), Hepa-1c1c7 cells (GSE154387), HPC7 cells (GSE48086), HSPC cells (GSE131583), kidney (GSE49847), liver (GSE91731), MEF cells (GSE99197), MEL cells (GSE181234), mature olfactory sensory neurons (GSE112153), nephron progenitor cells (GSE90016), anterior neural progenitor cells (GSE160654), pancreas (GSE59119), Patski cells (GSE59779), pro-B cells (GSE109909), round spermatids (GSE70764), small intestine (GSE49847), spinal motor neurons (GSE196170), spleen (GSE111772), splenic plasmablasts (GSE44637), stomach (GSE91488), testis (GSE49847), thymus (GSE180937), trophoblast stem cells (GSE110950). Human: A-673 cells (GSE185132), adrenal gland (GSE106071), ascending.aorta (GSE143103), astrocytes (GSE30263), B cell (GSE206145), bipolar neuron (GSE96269), 5637 cells (GSE193886), brain (GSE209256), brain microvascular endothelial cell (GSE30263), breast epithelium (GSE105608), CLB-Ga cells (GSE224242), CMP cells (GSE231486), colonic mucosa (GSE209074), coronary artery (GSE127477), DLD-1 cells (GSE142746), esophagus muscularis mucosa (GSE142970) , esophagus squamous epithelium (GSE105931), gastrocnemius medialis (GSE143020), gastroesophageal sphincter (GSE127521), GMP cells (GSE231486), Gp5d cells (GSE180150), ESC (GSE211101), ESC-derived SC-beta cells (GSE211101), HAP1 cells (GSE180691), HCT116 cells (GSE184106), heart right ventricle (GSE175279), HEK293T cells (GSE206145), HeLa S3 cells (GSE32883), HepG2 cells (GSE30226), HT1080 cells (GSE153869), Jurkat cells (GSE130140), K562 cells (GSE180175), Kidney (GSE33213), Kuramochi cells (GSE152885), L1207 cells (GSE193886), left lung (GSE188029), left ventricle myocardium inferior (GSE187266), liver (GSE127549), lower leg skin (GSE105391), limbal stem cells (GSE192625), lung (GSE33213), MCF 10A cell (GSE183381), MCF7 cell (GSE181460), MEP cells (GSE231486), mucosa descending colon (GSE208481), Mutu-1 cells (GSE160973), pancreas (GSE174993), Peyers patch (GSE105594), PLC/PRF/5 cells (GSE209849), 22Rv1 cells (GSE200168), psoas muscle (GSE209062), right atrium auricular (GSE127378), RPE cells (GSE196727), RT-112 cells (GSE193886), SD48 cells (GSE193886), SH-SY5Y cells (GSE141278), sigmoid colon (GSE105852), suprapubic skin (GSE139782), testis (GSE105739), thoracic aorta (GSE127422), thyroid gland (GSE105921), tibial artery (GSE105707), tibial nerve (GSE105554), U2OS cells (GSE175731), vagina (GSE105477).

